# Epigenetic aging waves: Artificial intelligence detects clustering of switch points in DNA methylation rate in defined sex-dependent age periods

**DOI:** 10.1101/2022.10.02.510495

**Authors:** Elad Segev, Tamar Shahal, Thomas Konstantinovsky, Yonit Marcus, Gabi Shefer, Yuval Ebenstein, Metsada Pasmanik-Chor, Naftali Stern

**Affiliations:** The Sagol Center for Epigenetics of Aging and Metabolism, Institute of Endocrinology, Metabolism and Hypertension, Tel Aviv-Sourasky Medical Center; Sackler Faculty of Medicine, Tel Aviv University, Israel; Department of Applied Mathematics, Holon Institute of Technology, Israel; Department of Engineering, Bar Ilan University, Israel; The Sackler Faculty of Medicine, Tel-Aviv University, Israel; Department of Chemistry, Tel Aviv University, Israel; Bioinformatics Unit, The George S. Wise Faculty of Life Science, Tel Aviv University, Israel

## Abstract

**Background:** Aging is linked to hypermethylation of CpG sites on promoters and enhancers, along with loss of methylation in intergenic zones. That such changes are not necessarily a continuous process is exemplified by the extensive changes in DNA methylation during development with another significant time of change during adolescence. However, the relation between age and DNA methylation during adult life has not been systematically evaluated. In particular, potential changes in methylation trends in the same CpGs over the years that may occur with aging remain largely unexplored.

**Methods:** Here we set out to determine the average trends by age of the CpG sites represented in the Illumina 450 platform, based on data from 2143 subjects of the age range of 20 to 80 years, compiled from 24 different cohorts. Using several mathematical procedures, we initially separated stationary probes from probes whose methylation changes with age. Among the latter, representing ∼20% of the probes, we then focused on the identification of CpG sites with switch points, i.e., a point where a stable trend of change in the age-averaged methylation is replaced by another linear trend.

**Results:** Using several mathematical modeling steps, we generated a machine learning model that identified 5175 CpG sites with switch points in age-related changes in the trend of methylation over the years. Switch points reflect acceleration, deceleration or change of direction of the alteration of methylation with age. The 5175 switch points were limited to 2813 genes in three waves, 80% of which were identical in men and women. A medium-size wave was seen in the early forties, succeeded by a dominant wave as of the late fifties, lasting up to 8 years each. Waves appeared∼4-5 years earlier in men. No switch points were detected on CpGs mapped to the X chromosome.

**Conclusion:** In non-stationary CpG sites, concomitant switch points in age related changes in methylations can be seen in a defined group of sites and genes, which cluster in 3 age- and sex-specific waves.

## Introduction

Age-related changes in DNA methylation have been recognized for more than four decades (1–3). In brief, global DNA methylation declines from the onset of adulthood to advanced age (4–7), but this trend is comprised of two opposing vectors: CpG sites with overall low DNA methylation, such as promoter-associated CpG islands, tend to increase methylation with age, whereas hypermethylated DNA zones, such as intergenic non-island CpGs tend to lose methylation with age. The resultant overall decrease in global DNA methylation with age reflects the fact that CpGs residing outside of CpG islands and tend to be hypermethylated outnumber the CpG sites in the CpG islands, whose methylation level rises with the passage of time (8-10). This also leads to a gradual shift in DNA methylation levels toward the mean with increasing age (8–12). Age-related changes in DNA methylation tend to occur preferentially at CpG island shores and shelves and enhancers (13). That certain CpG sites are closely and linearly related to age has been utilized by Horvath et al (4,5), Hannum (11), and Weidner et al (12) to formulate the concept of age/time related epigenetic clocks, according to which an epigenetic age, in years, can be mathematically derived from specific CpG sites such that it approximately parallels the chronological age. Shahal et al. (14) subsequently deconvoluted the Horvath’s epigenetic clock to its components, showing significant inter-individual variability. Furthermore, a large body of work now links upward drifting from the epigenetic age, which reflects accelerated epigenetic aging, to earlier mortality, decreased healthy longevity and a number of diseases, such as breast cancer (15–21). Since most epigenome-wide DNA methylation data is now based on large platforms identifying the level of methylation in predefined CpG sites, it is of interest that Florath et al (7) who analyzed >480000 CpG sites in whole blood DNA of a population-based cohort study aged between 50 and 75 years, found only 162 CpG sites with the high Spearman correlation coefficients (R>0.6) between DNA methylation and age.

Whereas these and likely other CpG sites selected to comprise the basis for “epigenetic clocks” apparently serve as excellent aging markers, or may even be causatively linked to the aging process, they are obviously too few to account for the large age-related changes seen in DNA methylation. Because many reports revealed the linear relation to age and its link to health conditions, we sought, in the present report, to explore the overall pattern/s of changes in DNA methylation as a function of age. However, non-liner changes in DNA methylation with age have been previously reported in humans (22) and canids (23). Extreme examples of large changes in DNA methylation are known to occur in specific ages in early life: after birth, average DNA methylation levels increase in blood throughout the first year of life (24,25). Likewise, during adolescent transition changes in DNA methylation were observed in more than 15000 CpGs, many of which were associated with genes relevant to cell growth and immune system development (26).

In the present study we pursued the following aims: 1) To identify non-stationary CpG sites, i.e., all CpG methylation sites that undergo detectable age-related changes between the age of 20 and 80 years. 2) To identify CpG sites with age-related switch point (SP), i.e., CpG sites whose methylation fraction changes in a curvilinear manner such that changes in DNA methylation are either accelerated, decelerated or switch their trend direction at a certain age. 3) To investigate the possibility that acceleration or deceleration in methylation cluster around defined age-related zones (e.g., perimenopause). To this aim we used a compiled data base constructed of 24 different published cohorts who all used the Illumina 450K human DNA methylation platform.

## Methods

We extracted data from published databases and performed several analytical procedures resulting in the development of an artificial model that can identify the CpGs with a SP (epigenetic switch point; ESP) in their mean methylation fraction between the ages of 20 and 80 years. The entire process is described in Fig1.

**Figure 1:**
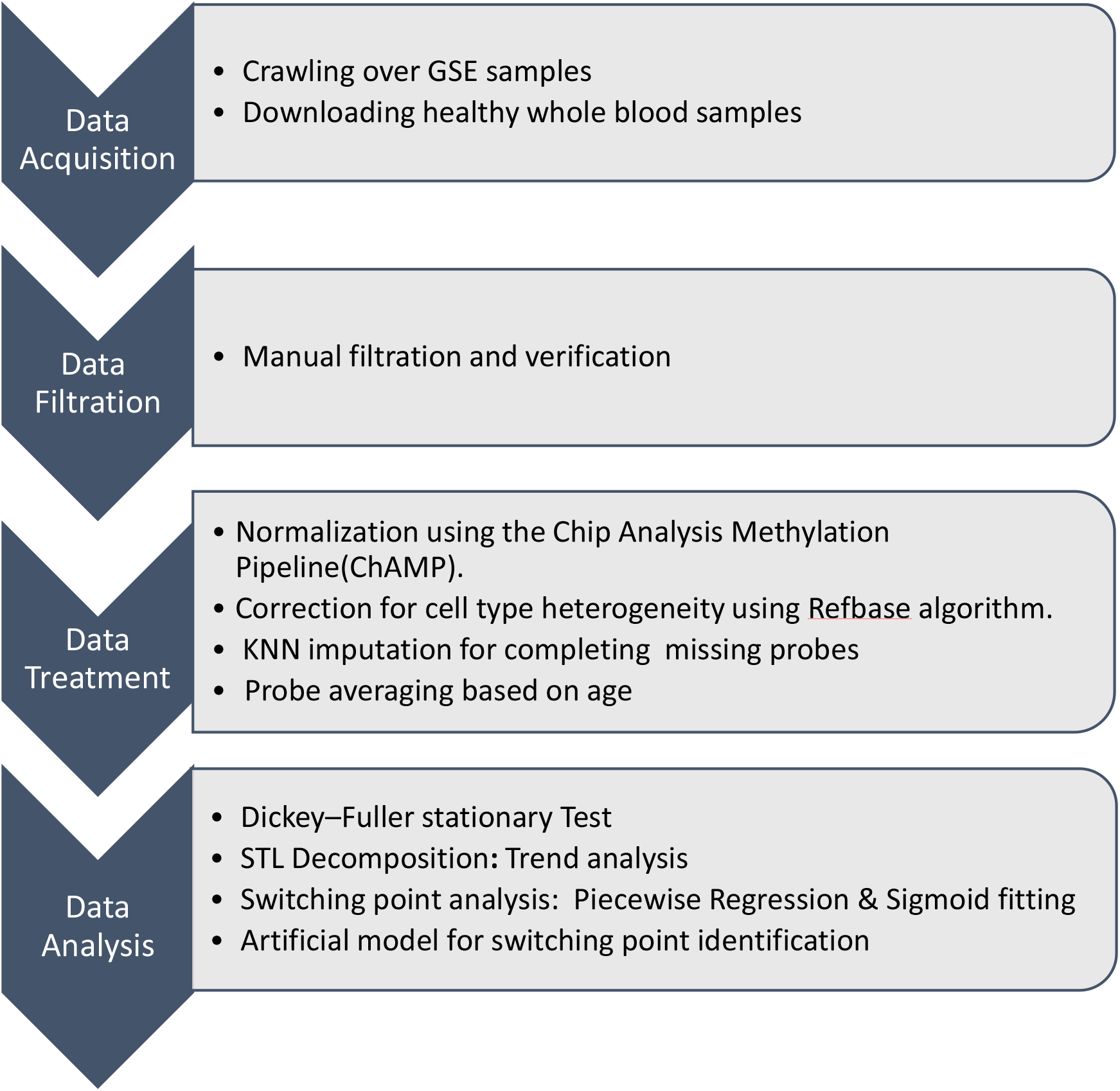
A schematic overview of the data extraction and analytical procedures used for the identification of ESP.

### Data acquisition and processing

#### 1. Crawling over GSE samples

We used an extensive automatic web crawler to search for all available repositories that match our query for whole blood and healthy subjects at the age range of 20–80 years (14) from the Gene Expression Omnibus (GEO) Data sets repository.

#### 2. Downloading healthy whole blood samples

All relevant β values and idat values were downloaded, Samples with idat values were converted to β values using champ package (27)

#### 3. Data Filtration and verification

All samples were manually inspected. Samples from datasets that were not labeled as healthy and contained case-control groups were manually examined to ensure that control samples were indeed healthy. The entire process yielded a total of 2221 samples from 24 different data sets, which were then subjected to further analysis

#### 4. Data treatment

The β values of all filtered samples were normalized using BMIQ code “champ.norm” implemented at Champ package (28).

#### 5. Correction for cell type heterogeneity

We corrected for cell type heterogeneity using “champ.refbase” implemented at Champ package (29).

#### 6. K-nearest neighbors’ algorithm [KNN] for completing 47 missing probes

To maximize the preservation of CpG sites and remove poor-quality samples across the different datasets we used a k-nearest neighbors’ algorithm (KNN) based approach. To infer the degree of relative similarity between the different samples based on their CpG site β values, the sample-linked gender, age and the percentage of missing CpG sites were entered. In the process of searching for the nearest neighbors sharing the same metadata (age, gender, similar β values) and emplacing the missing β values with the values of the closest neighbor, 47 samples could be thus adjusted and were retained. However, 78 other samples for which no nearest neighbor were removed from our dataset. The entire process yielded a total of 2221 samples from 24 different data sets, which were then subjected to further analysis (table 1S). Of the resulting 2143 samples, 1119 were females and 1024 males, with an age distribution presented in figure S1, supplementary file S1.

**Table 1:** Age related significantly enriched Kyoto Encyclopedia of Genes and Genomes (KEGG) biological pathways in the SP genes list

| Pathway | # of Genes | Fold Enr* | Benjamini | Age related changes |
| --- | --- | --- | --- | --- |
| hsa04261:<br>Adrenergic signaling in cardiomyocytes | 47 | 2.3 | 1.5E-05 | Decline in heart performance and myocardial remodeling ability; decrease in cardiomyocytes survival ability, which involves reduced response to adrenergic signaling |
| hsa04360:<br>Axon guidance | 52 | 2.1 | 2.3E-05 | Dysregulation of axonal mRNA localization and local protein translation |
| hsa04713:<br>Circadian entrainment | 32 | 2.4 | 1.3E-04 | Increasingly dysregulated with age; disruption of circadian entrainment accelerates aging |
| hsa04725:<br>Cholinergic synapse | 34 | 2.2 | 3.2E-04 | Cholinergic synapse function declines in the brain and neuromuscular junction. |
| hsa04611:<br>Platelet activation | 36 | 2.1 | 3.6E-04 | Increases with age → increased cardiovascular thromboembolic clinical and subclinical events; noncanonical platelet effects accelerate neurological diseases including Alzheimer's disease, possibly by increased secretion of beta-amyloid. |
| hsa04911:<br>Insulin secretion | 27 | 2.3 | 1.0E-03 | Aging beta cells display lower insulin secretory capacity in response to cellular and circulating cues |
| hsa04915:<br>Estrogen<br>signaling<br>pathway | 36 | 1.9 | 2.8E-03 | Beside traditional tissue targets for estrogens (bone and cardiovascular system), decline in estrogen activity unfavorably affect the preservation of brain connectivity, synapse structure and function |
| hsa04930:<br>Type II<br>diabetes<br>mellitus | 15 | 2.4 | 1.9E-02 | Aging is modulated by metabolic dysregulation of carbohydrate, fat and protein metabolism, overlapping with or culminating type 2 diabetes |
| hsa04211:<br>Longevity<br>regulating<br>pathway | 22 | 1.8 | 4.4E-02 | Normal insulin response, caloric restriction, maintenance of autophagic flow and sufficient mitochondrial biogenesis and controlling inflammation and oxidative stress, improve survival and longevity and retard aging |

\*Fold enrichment; # = number

#### 7. Probe averaging based on age

Probes missing in more than 1% of our dataset were filtered out, leaving 341,247 probes. The 2143 samples were tabulated by age and the average β for each CpG site was calculated for each age (in years, between 20-80). This data structure allowed us to remove noise and converge to age-specific trends. Moreover, it reduced the data spread from 2143 columns for each of the 341,247 probs to 61 columns, a ∼35-fold condensation which allowed more extensive calculations. Hence, we ended up handling a data set of 61 columns, one column for each year of age (20-80 years) with a mean β value for each CpG site in each of the cells. This data representation not only reduced the size of the data analyzed but also can be viewed as a signal sampled with a frequency of 1 year, which allows the usage of a vast variety of tools, algorithms and mathematical models derived from the branches of time series analysis to uncover hidden aging patterns that may appear as methylation β values. An example as to how such a signal of a CpG site looks before and after the averaging can be seen in Figure 3A, B, E, F.

#### 8. Dickey–Fuller stationary Test

Since we strived to focus on CpG sites that undergo significant change over the age years of 20-80, we applied the augmented Dickey–Fuller test, which afforded the identification of stationary probes, i.e., probes that do not change significantly (p <0.05) in terms of their means, standard deviations (SD) as well as seasonal (cyclic, sinus-like) behavior. This is conceptually depicted in Figure 2, in which 4 hypothetical probes are presented. The black curve represents a stationary probe which undergoes no significant change in the beta value or SD. Such hypothetical signal probes can be statistically described as discrete variables with normal distribution and a given mean and SD. Since there is no significant change across the years, in either the beta value or the SD, these probes were not further considered for evaluation in the present report.

**Figure 2:**
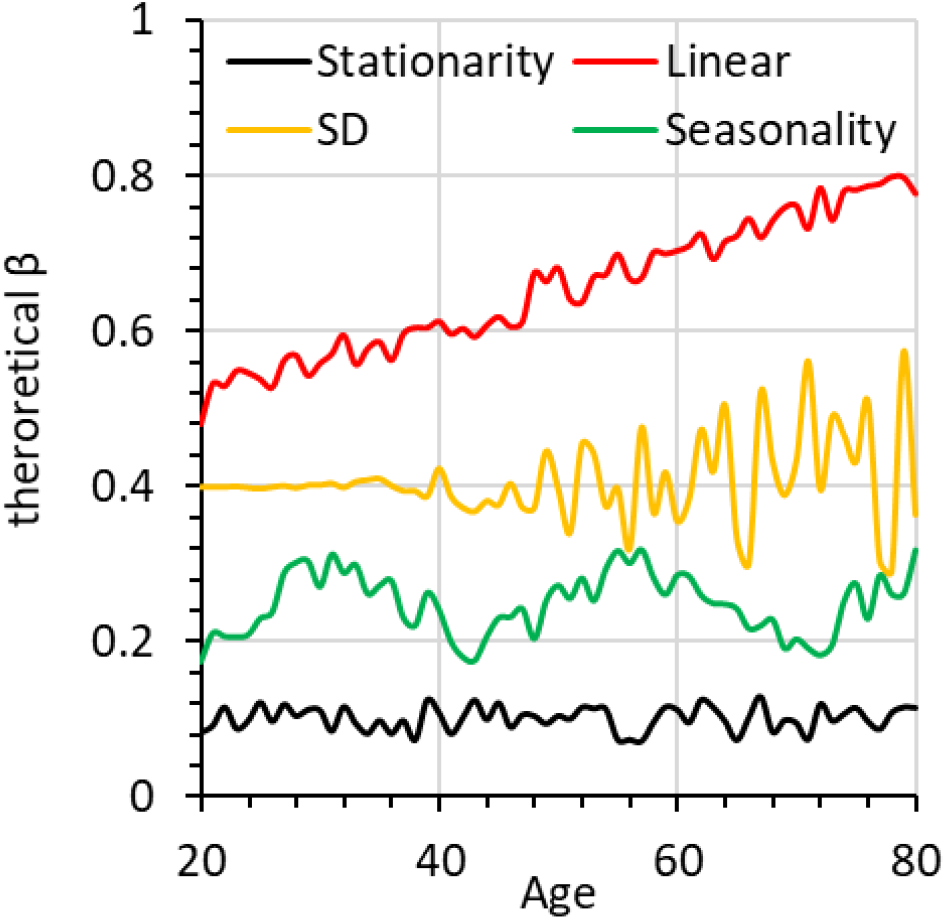
Four hypothetical CpG signals: in black, a stationary signal; the red, orange and green curves exemplify three different non-stationary signals: a signal with an ascending change in beta value over time in red; in orange, a signal showing a change in SD value over time, and in green a signal with seasonal behavior, i.e., a sine like behavior.

On the other hand, the red orange and green curves are non-stationary signals representing change in beta value, SD, or seasonality behavior (sine function-like behavior), respectively, over time. In all, this analysis revealed 69,275 “non-stationary” CpG cites.

#### 9. STL Decomposition / Trend analysis

Seasonal and Trend using Loess decomposition (STL) was used to reduce noise (30). This mathematical method allows the splitting of the CpG signal over time into three components: trend, seasonal and a remainder (noise) component. The average methylation curve between 20 to 80 years before and after STL process can be seen for probes cg03037684 and cg16803083 mapped to ESR1 gene and to a region that was not annotated, respectively in figure 3B and C, and in figure 3 F and G, respectively.

**Figure 3:**
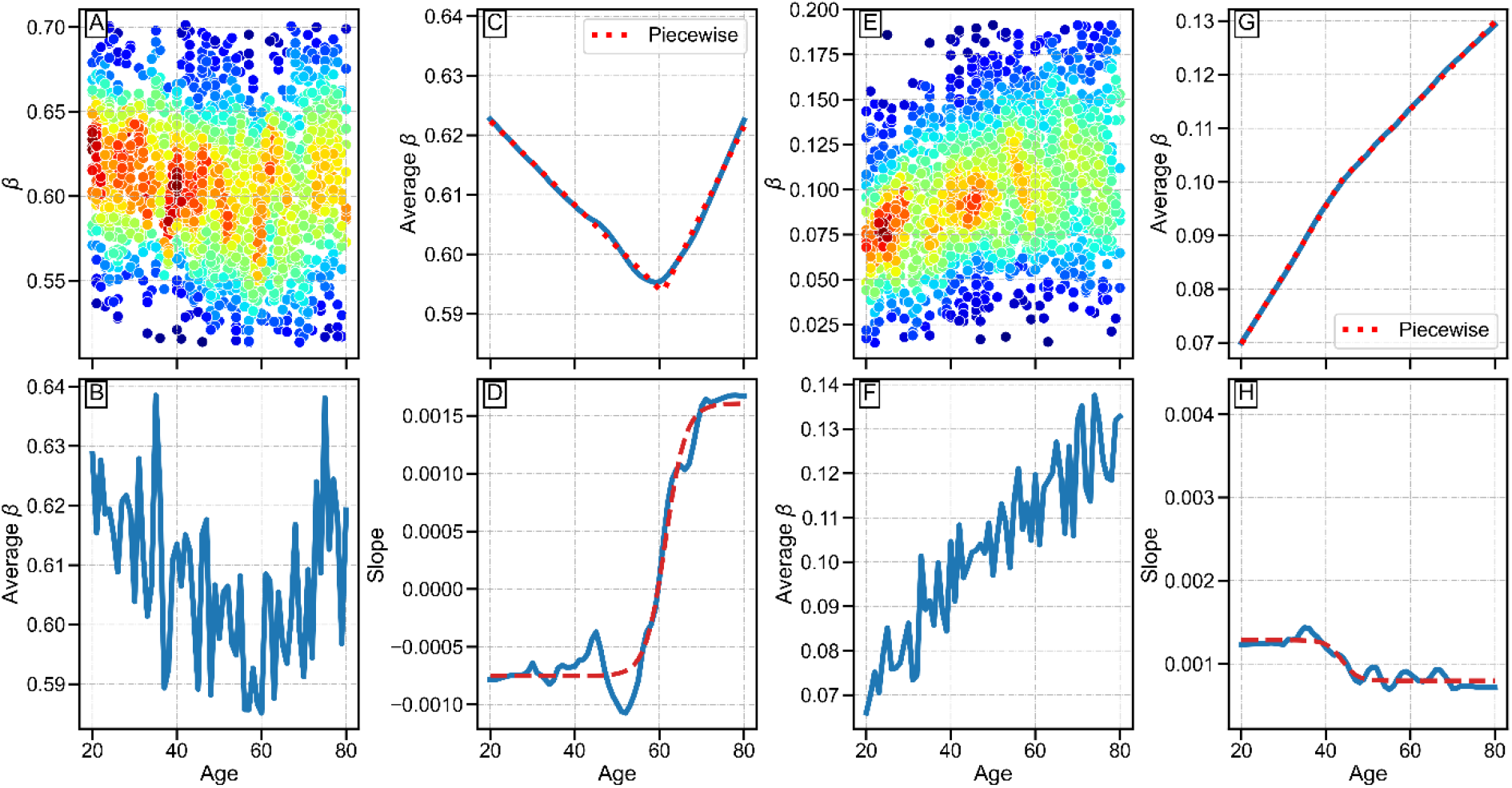
Mean change in β values of two CpG sites between the ages of 20 to 80 years. A-D and E-H represent 2 CpG sites with a change in mean methylation fraction with age, one with (cg03037684; mapped to ESR1; curvilinear change) and the other without an ESP (cg16803083; linear change), respectively. (A, E) the original data of the β values of 2143 individuals between 20 to 80 years old, (B, F) Average β values of all individuals (presented in A, E, respectively) at the same age with one year resolution, (C, G) An STL analysis of the average values presented in B and F (blue line), a piecewise model fitting to the STL data is presented as two linear red dashed lines, and (D, H) The first derivative of the averaged data following SLT, yields a sigmoid shape curve in D and a close to horizontal shaped curve in H (blue lines). The dashed red line shows the close fitting of a sigmoid to the STL average data. Color code in A and E is for the density of the number of individuals for each beta value and for each age (per one year interval; red for most dense and blue for list dense).

#### 10. Switch point analysis-Piecewise Regression & Sigmoid fitting

The previous filtering eliminated CpG sites showing no significant change with age. We next identified CpG sites with ESP, i.e., CpG sites having a definable behavior until a certain age, but showing a change in behavior as of a specific age and on as demonstrated in figure 3A-D. To this aim we combined two methods which generated parameters for the construction of an artificial model that we subsequently used for the identification and selection of ESPs.

A. *Piecewise regression analysis*, which is based on fitting two linear curves to the average methylation values at each year of age between 20 to 80 years (red dashed lines in figure 3C and 3G) as formulated in equation 1, where (*α*_1,_ *α*_2,_ *β*_1,_ *β*_2_) are the intercepts and slopes of each of the linear curve, respectively, calculated based on the least squares, and *S*_*p*_ is the switch point.

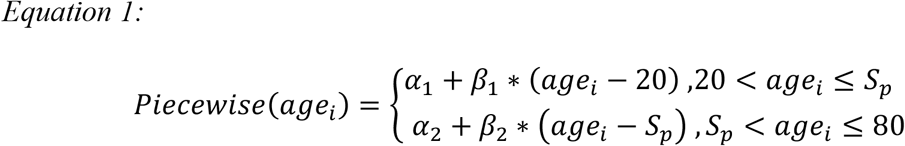 The age of switch point is where the two fitted linear curves meet.
B. *Sigmoid fitting analysis*, which is based on fitting a sigmoid curve to the first derivative of the average β values, between 20 to 80 years, with 1year intervals for each of our non-stationary CpGs sites. In Figure 3D and 3H the first derivatives of average β values are presented as blue lines and the fitting to a sigmoid as dashed red lines, as described in equation 2, where x is the age, *λ*the middle of the sigmoid and S,B, and K are scaling parameters (supplementary file S1).

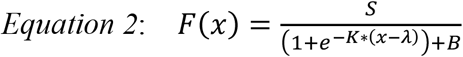 The least squares method was used to calculate the optimal parameters (S, B, K, λ). The age of SP, then, is the inflection point of the sigmoid curve (indicated by the vertical grey dashed arrow in Figure 3D).

The piecewise analysis provides a very ideal behavior of a particular time point where the linearity of a beta values of a CpG site changes. Mathematically it is equivalent to a single point in which the slope changes and can be defined as discontinuous for its first derivative (Figure 4A). In population studies, a switch point will not be seen insentiently at a certain age but will rather take place gradually over several years. Therefore, a sigmoid curve, fitted to the first derivative of our non-stationary, STL-processed average β values between 20 to 80 years, may better describe the gradual emergence of a SP, as illustrated in figure 4B). According to this analysis, the behavior of ESPs can be also characterized by a bilinear course having the first slope of methylation loss/ gain rate up to a certain age range and a second different slope from that age range onwards, as described in figure 3C. This is depicted by the two green arrows, where the vertical green lines denote the start and the end of an ESP period.

**Figure 4:**
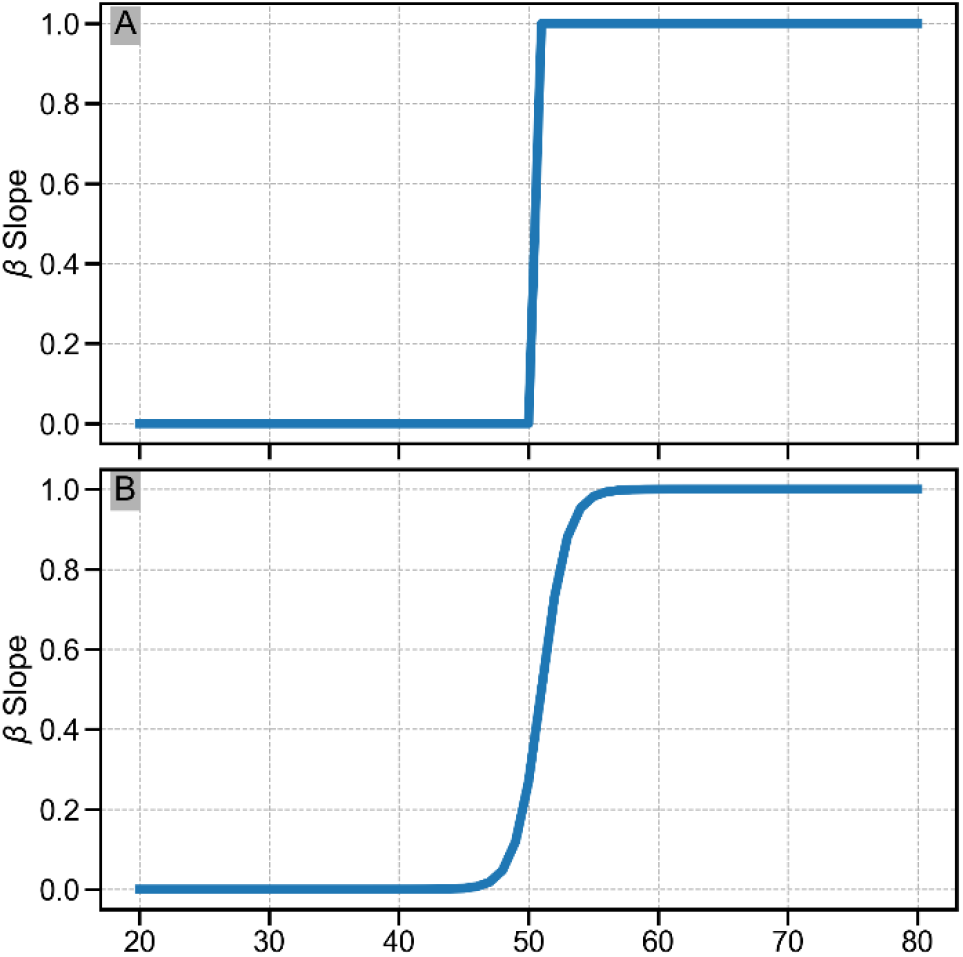
illustration of the first derivative of β values of a SP CpG site from 20 to 80 years. (A) an instantaneous SP, (B) a gradual, more realistic SP.

#### 11. Artificial model for the identification of switch points

To filter out poor quality and/or noisy probes, we constructed a mathematical model resulting in a collection of parameters which were used to distinguish between valid and non-valid SP CpGs. The model was based on an in person, CpG by CpG, visual inspection of randomly selected 1500 CpG sites and their features (as detailed below) out of the entire non-stationary 69,275 CpG sites population. During this procedure each screened CpG was labelled as having/not having a SP. ESP sites were identified as such only if any of the following exclusion criteria was present: a) the first derivative of the curve of the average β values as a function of age, did not fit a sigmoid curve, as indicated, for example, by the horizontal, first derivative curve derived from a linear methylation gain with age (figure 3H); b) derived STL was incompatible with the original data curve, c) the STL suggested the existence of a SP, but the actual curve could be fitted into an alternative pattern; d) a SP point generated by an abrupt and large change in beta values, which is likely fortuitous secondary to technicalities as may be suggested from the illustration in figure 4A. The piecewise regression and sigmoid models were used to derive 19 parameters (table 2S in supplementary). Based on in person visual inspection of the 1500 CpG sites, a decision tree model was trained and evaluated with an average F1 Score of 0.89 on 5 fold cross validation. The decision tree model was then used to examine all 69,275 non-stationary probes in order to distinguish valid ESP probes from invalid probes. Eventually, 5175 probes that passed this filtration process were identified as ESPs.

**Table 2:** Prominent aging and longevity related genes associated with SP CpG sites

| SP- aging<br>gene<br>Symbol | SP- Aging genes | probe | UCSC<br>RefGene<br>Group | Overall<br>slope<br>trend | Mean<br>$\beta$ | SP<br>age |
| --- | --- | --- | --- | --- | --- | --- |
| ADCY5 | Adenylate Cyclase 5 | cg25661931 | 1stExon | + | 0.97 | 61 |
| ADIPOQ | Adiponectin, C1Q And Collagen<br>Domain Containing | cg10681525 | Body | + | 0.71 | 55 |
| ADRA1B | Adrenoceptor Alpha 1B | cg09444685 | Body | - | 0.68 | 64 |
| AGER | Advanced Glycosylation End-Product Specific Receptor | cg10996463 | Body | - | 0.87 | 43 |
|  |  | cg11105830 | Body | + | 0.81 | 45 |
|  |  | cg18139800 | Body | + | 0.92 | 58 |
| ALOX15B | Arachidonate 15-Lipoxygenase<br>Type B | cg15799267 | 5'UTR | - | 0.34 | 55 |
| APOB | Apolipoprotein B | cg13287979 | Body | + | 0.83 | 55 |
|  |  | cg03350299 | TSS200 | + | 0.43 | 56 |
|  |  | cg25071744 | TSS200 | + | 0.52 | 57 |
|  |  | cg23949611 | Body | - | 0.83 | 59 |
| ATG7 | Autophagy Related 7 | cg01796438 | TSS1500 | + | 0.42 | 55 |
|  |  | cg11277834 | 5'UTR | + | 0.69 | 61 |
| BCL2 | BCL2 Apoptosis Regulator | cg08223235 | Body | + | 0.31 | 57 |
| BNIP3 | BCL2 Interacting Protein 3 | cg15390444 | Body | + | 0.69 | 55 |
| CLDN1 | Claudin 1 | cg03601836 | Body | - | 0.87 | 67 |
| CREBBP | CREB Binding Protein | cg04141008 | Body | + | 0.72 | 55 |
| CSNK1E | Casein Kinase 1 Epsilon | cg22494725 | Body | - | 0.63 | 65 |
| DUSP1 | Dual Specificity Phosphatase 1 | cg12333707 | TSS1500 | + | 0.50 | 60 |
| EGFR | Epidermal Growth Factor<br>Receptor | cg05246100 | Body | - | 0.72 | 59 |
|  |  | cg16488565 | Body | + | 0.55 | 57 |
|  |  | cg16751451 | TSS1500 | + | 0.51 | 60 |
| ERCC1 | ERCC Excision Repair 1,<br>Endonuclease Non-Catalytic<br>Subunit | cg23347323 | 3'UTR | + | 0.73 | 55 |
| ESR1 | Estrogen Receptor 1 | cg03037684 | 3'UTR | - | 0.61 | 58 |
| ETS1 | ETS Proto-Oncogene 1,<br>Transcription Factor | cg01900413 | Body | + | 0.25 | 62 |
|  |  | cg25068347 | Body | + | 0.77 | 66 |
|  |  | cg21695395 | Body | + | 0.60 | 67 |
| ETS2 | ETS Proto-Oncogene 2,<br>Transcription Factor | cg15892280 | 5'UTR | - | 0.43 | 55 |
| FOXO3 | Forkhead Box O3 | cg08792630 | Body | + | 0.37 | 63 |
| HDAC4 | Histone Deacetylase 4 | cg14653043 | Body | + | 0.93 | 48 |
|  |  | cg24401044 | Body | - | 0.97 | 51 |
|  |  | cg01771737 | Body | - | 0.79 | 52 |
|  |  | cg04438064 | Body | - | 0.49 | 53 |
|  |  | cg05758467 | Body | + | 0.45 | 57 |
|  |  | cg15002163 | Body | + | 0.61 | 61 |
|  |  | cg11550064 | Body | + | 0.61 | 62 |
| HIPK2 | Homeodomain Interacting<br>Protein Kinase 2 | cg03520802 | Body | - | 0.54 | 53 |
|  |  | cg00995324 | Body | + | 0.92 | 61 |
|  |  | cg23646343 | Body | + | 0.25 | 68 |
| HK3 | Hexokinase 3 | cg19791262 | TSS1500 | + | 0.73 | 51 |
|  |  | cg14709481 | 5'UTR | + | 0.68 | 56 |
| IGF1R | Insulin Like Growth Factor 1<br>Receptor | cg08920032 | Body | - | 0.49 | 58 |
| IGF2R | Insulin Like Growth Factor 2<br>Receptor | cg21774926 | TSS1500 | + | 0.37 | 59 |
| IGFBP2 | Insulin Like Growth Factor Binding Protein 2 | cg20366479 | Body | + | 0.56 | 44 |
| INSR | Insulin Receptor | cg23075968 | Body | + | 0.39 | 61 |
| MGST1 | Microsomal Glutathione S-Transferase 1 | cg22494907 | 5'UTR | - | 0.91 | 61 |
| MTOR | Mechanistic Target Of Rapamycin Kinase | cg10315903 | TSS1500 | + | 0.65 | 62 |
| NCOR2 | Nuclear Receptor Corepressor 2 | cg13146040 | 5'UTR | + | 0.88 | 46 |
|  |  | cg03821418 | Body | + | 0.54 | 52 |
|  |  | cg07241090 | Body | + | 0.35 | 56 |
|  |  | cg22820108 | 5'UTR | - | 0.40 | 57 |
|  |  | cg17825194 | Body | - | 0.34 | 57 |
|  |  | cg22700848 | 5'UTR | + | 0.38 | 58 |
|  |  | cg17187521 | 5'UTR | + | 0.40 | 60 |
|  |  | cg17387577 | Body | + | 0.33 | 69 |
| NEIL1 | Nei Like DNA Glycosylase 1 | cg02426940 | Body | + | 0.82 | 56 |
| NEK6 | NIMA Related Kinase 6 | cg13958199 | TSS1500 | + | 0.61 | 53 |
|  |  | cg14082739 | 5'UTR | + | 0.84 | 49 |
| NFE2L1 | NFE2 Like BZIP Transcription Factor 1 | cg27568306 | Body | + | 0.91 | 53 |
| NGF | Nerve Growth Factor | cg17750109 | 5'UTR | - | 0.80 | 49 |
|  |  | cg02987481 | TSS1500 | + | 0.43 | 59 |
| NOS1 | Nitric Oxide Synthase 1 | cg21006686 | TSS1500 | + | 0.70 | 57 |
|  |  | cg02500231 | 5'UTR | + | 0.25 | 71 |
| NOS3 | Nitric Oxide Synthase 3 | cg00571021 | Body | - | 0.00 | 0 |
| NOTCH3 | Notch Receptor 3 | cg08529654 | Body | + | 0.41 | 60 |
| PAX4 | Paired Box 4 | cg11975652 | 3'UTR | + | 0.68 | 49 |
| PIK3CD | Phosphatidylinositol-4,5-Bisphosphate 3-Kinase Catalytic Subunit Delta | cg23251761 | 5'UTR | - | 0.73 | 49 |
|  |  | cg19267205 | Body | + | 0.41 | 56 |
| SLC6A6 | Solute Carrier Family 6 Member 6 | cg07573937 | 5'UTR | - | 0.51 | 31 |
| SLC9A3R2 | SLC9A3 Regulator 2 | cg09462956 | Body | + | 0.56 | 56 |
| TNFSF13 | TNF Superfamily Member 13 | cg13358186 | 5'UTR | - | 0.00 | 0 |
| TP53 | Tumor Protein P53 | cg07760161 | 5'UTR | + | 0.67 | 61 |
| TXNIP | Thioredoxin Interacting Protein | cg19389852 | 1stExon | + | 0.46 | 60 |
| VWF | Transmembrane Protein 270 | cg09117673 | TSS200 | - | 0.79 | 55 |
| WT1 | WW And C2 Domain Containing 1 | cg15946571 | Body | + | 0.94 | 59 |

### Pathway enrichment analysis for ESP genes

Kyoto Encyclopedia of Genes and Genomes (KEGG) enrichment analysis (31) was performed using DAVID (32) based on the 2,813 genes mapped to the 5175 ESPs. Pathways with Benjamini < 0.05 were considered as significantly enriched.

### Presence of ESP in genes associated with aging and longevity

We compiled a list of aging and longevity genes by assembling genes from four databases that are gathered in The Human Ageing Genomic Resources (HAGR) https://genomics.senescence.info/ (33): 1) GenAge: Database of Aging-Related Genes, which includes a curated database of over 300 genes related to <u>human aging</u> and a database of over 2000 aging- and longevity-associated genes in <u>model organisms</u> (34,35), 2) GenDR: Database of Dietary Restriction-Related Genes based on genetic manipulation experiments and gene expression profiling (36,37), 3) LongevityMap: a database of human genetic variants associated with longevity of over 2000 genes, some were found to be essential and some had no relation to longevity, 4) CellAge: Database of Cell Senescence Genes (38). Only genes that were found in either human, mouse or human cultured cells, were included in our list. From the LongevityMap database, we only included the genes that were found essential to longevity and deleted the non-essential ones. Finally, we deleted duplications and came up with a list of 973 resource genes related to aging and longevity. Venny 2.0.2 (39) was used to check for common ESP genes and aging/ longevity related genes.

## Results

### Stationary vs. non-stationary CpG sites: change in DNA methylation level in relation to age and the identification of CpG sites with an ESP behavior

We analyzed the trends of the average methylation level (β values), calculated for each year of age at each age point, from the age of 20 to the age of 80 years and found that the vast majority of the ∼340K CpG sites (n=341,247), eventually available for analysis, were stationary whereas 69,275 were found to be “non-stationary” (Materials and methods). We next examined the behavior of the non-stationary probe population and selected these with a clear ESP. The age of ESPs was the time at which acceleration/deceleration of DNA methylation gain/ loss rate occurred or the time at which a constant loss/ gain changed to a constant gain/ loss in methylation, respectably. In all, we identified a total of 5175 ESPs, of which 5113 were present in men and 4067 for women. Most were common for both sexes (supplementary file 2).

### ESP patterns

DNA methylation trends with ESPs over the age range of 20 to 80 years appeared at several different patterns. First, the overall trend of methylation change either rose or declined with age (green arrows in the positive and negative overall trends in figure 5A). The overall trend in the fraction of methylation for each of these CpGs followed one of five sub-SP patterns, based on the relative relation between the first and second slopes of methylation change rate, before and after the ESP period (figure 5A). There were more ESPs with an overall positive methylation than negative methylation trend, especially in men. Pattern III, characterized by a decrease in methylation up to a certain age followed by an increase in methylation from the age period of ESP onwards, was the most dominant ESP pattern for both men and women, in both overall trends (2465 and 1290 for positive and negative overall trends, respectively out of 5113 men ESPs and 2168 and 1676 for positive and negative overall trends, respectively, out of 4067 women ESPs). A sizable number of 1002 ESPs, showed acceleration in DNA methylation increase rate with age (pattern V), and this was seen exclusively in men.

**Figure 5:**
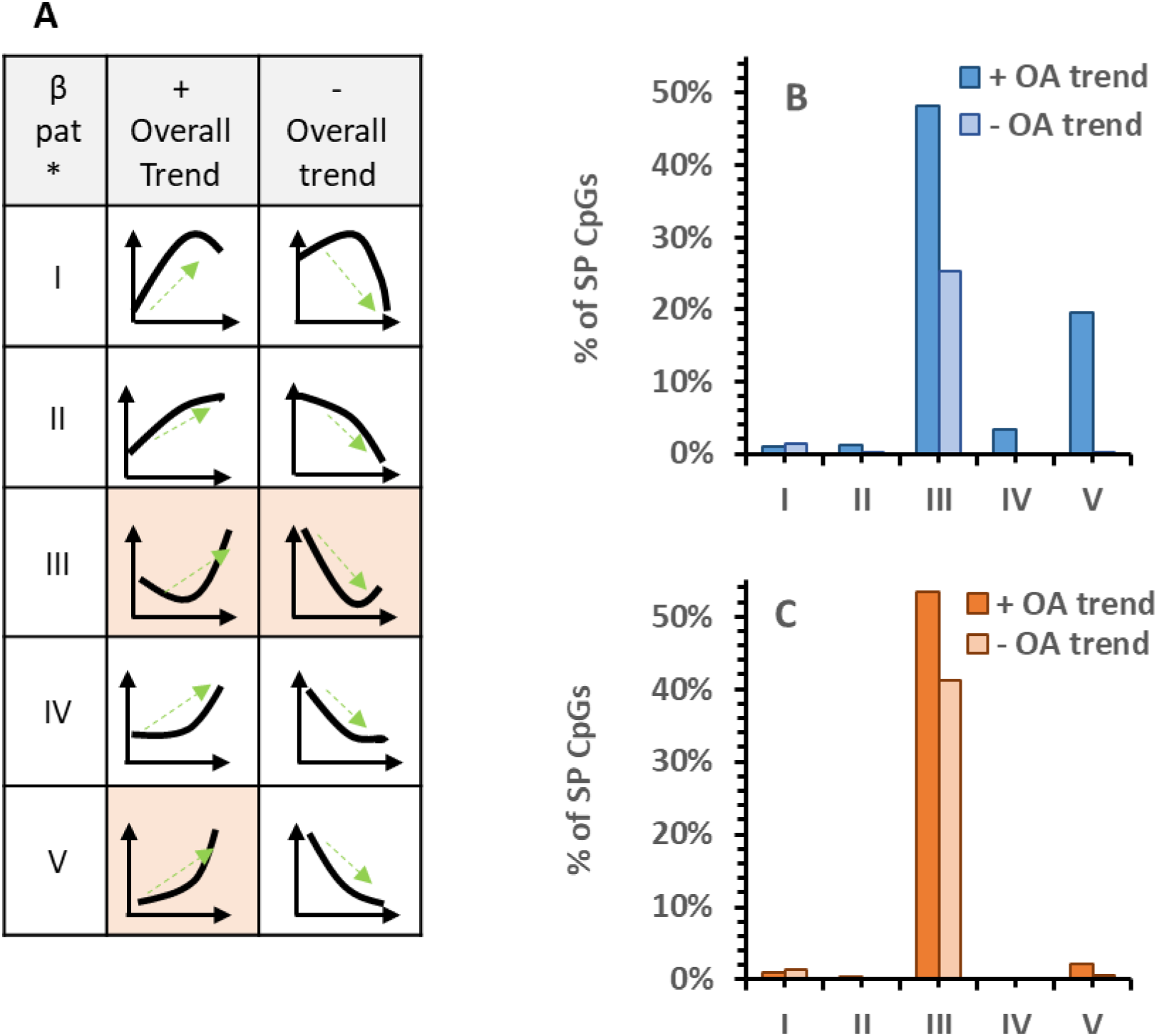
(A) a schematic view of the possible methylation SP patterns with age, for positive (+) and negative (-) overall trends (OA). Shown is the frequency of the different SP patterns in men (blue) (B) and women (orange) (C). SP CpG sites, for both positive and negative overall trends (green arrows in A), where 100% is the total SP CpG sites for each gender.

There was a relative even distribution of the mean beta value of the CpG probes with ESPs between 0.3 to 1, in CpGs with both negative and positive overall slop, for both men and women (supplementary file 1, figure S2). This is in complete contrast to a typical beta value distribution of the total 450K probes presenting CpGs in which most of the probes have either a beta value close to 0.2 or to 0.9 (28). This means that compared to the total CpGs population, a higher fraction ESPs have beta values around 0.5, which may imply a more dynamic behavior between the methylated and de-methylated state. It is also possible that more ESPs are mapped to imprinted genes. None of the CpGs resided on the X chromosome and hence, the higher beta values in probes with ESP are not due to x-inactivation.

### The age at which the epigenetic waves occur: relation to sex

Each ESP occurs at a definable age. Figure 6 demonstrates the frequencies of SP ages for all ESPs, separately for men and women (5113 and 4067 ESPs for men and women, respectively). The earliest and smallest (n=115, 98; M/F, respectively) wave of ESPs lays between the ages of 30 to 33 years for both genders. A second wave, which includes several hundred ESPs (n=765, 445; M/F, respectively) is seen between the ages of 45-51 years and 42-48 years for men and women, respectively. This ESP age period for women begins slightly earlier than that of men. Finally, the largest and dominant wave of SPs is discernible between the ages of 54-62 years and 60-69 years in men and women (n=4234, 3524; M/F respectively), respectively. Hence, most SP tend to appear later in women than in men. Indeed, 2 out of the 3 waves, and collectively, the vast majority of ESPs take place ∼six years and end ∼seven years later in women compared to men).

**Figure 6:**
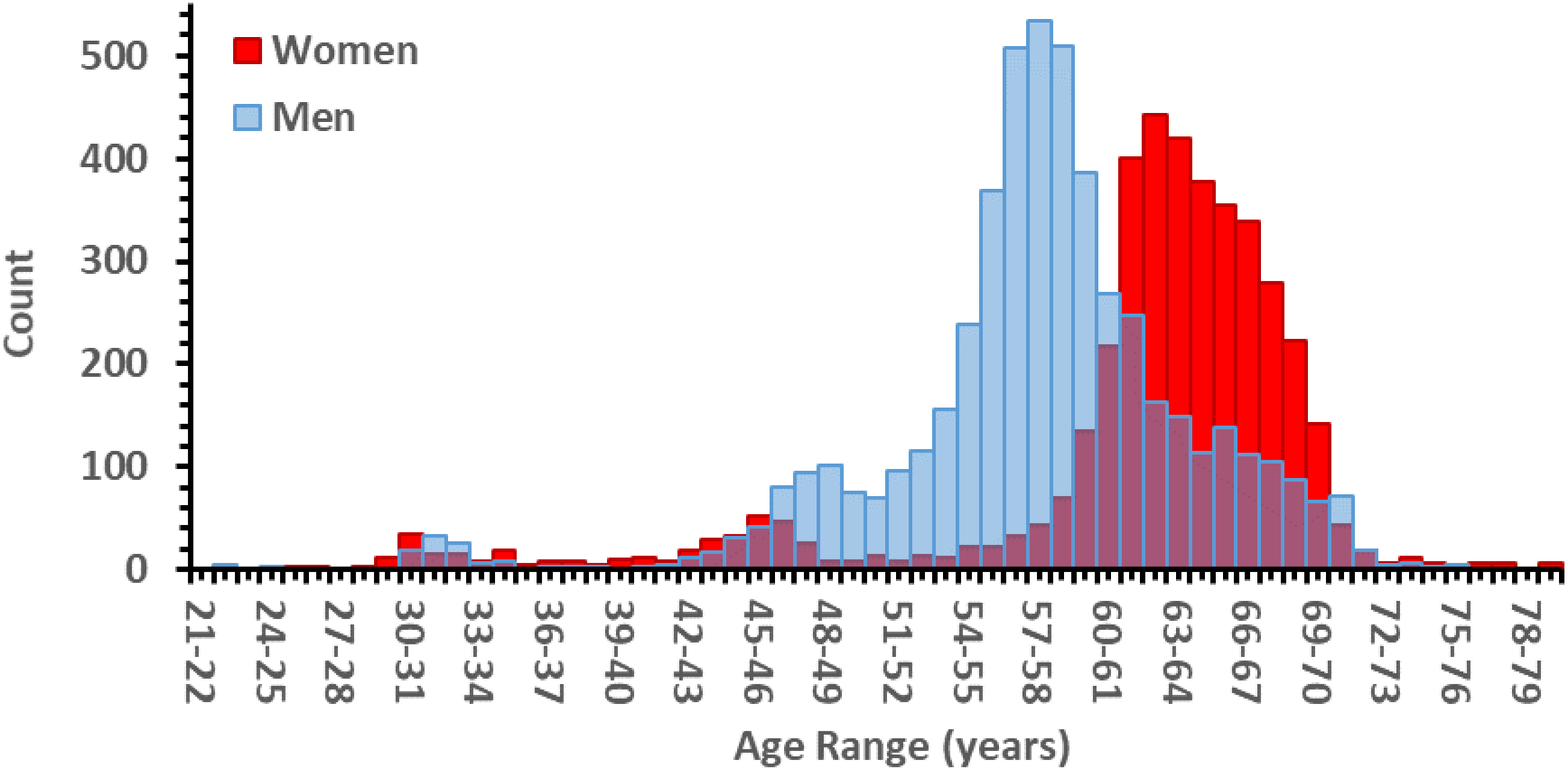
Sexual dimorphism in DNA methylation switch points as a function of age: Bimodal sex related distribution. The amount of CpGs with a switch point at a specific age, at 1 year intervals.

### Most SP CpG are common for women and men

Figure 7A is a Vann diagram showing that most ESPs are common for men and women (77.4%) while 21.4% ESPs are unique for men and only 1.5% are unique for women. We next constructed a Vann diagram for the 2,813 genes mapped to the ESPs (Figure 7B). In parallel to the ESPs, most genes with ESPs (81.4%) are common for men and women, 17.7% are unique for men and 0.9% are unique for women. This may imply that the aging process of men and women is mostly linked to similar biological pathways, with a clear difference in the age of onset. The full list of 5175 ESPs, their beta values, overall trends, first and second slopes (before and after the ESP period), age at the specific ESP, their mapped genes and their location in relation to their related gene, is listed in supplementary file S2. Since more than one CpG could be mapped for the same gene and since some CpGs were not mapped to a specific gene, the number of genes with ESPs is smaller than that of SP CpG sites.

**Figure 7:**
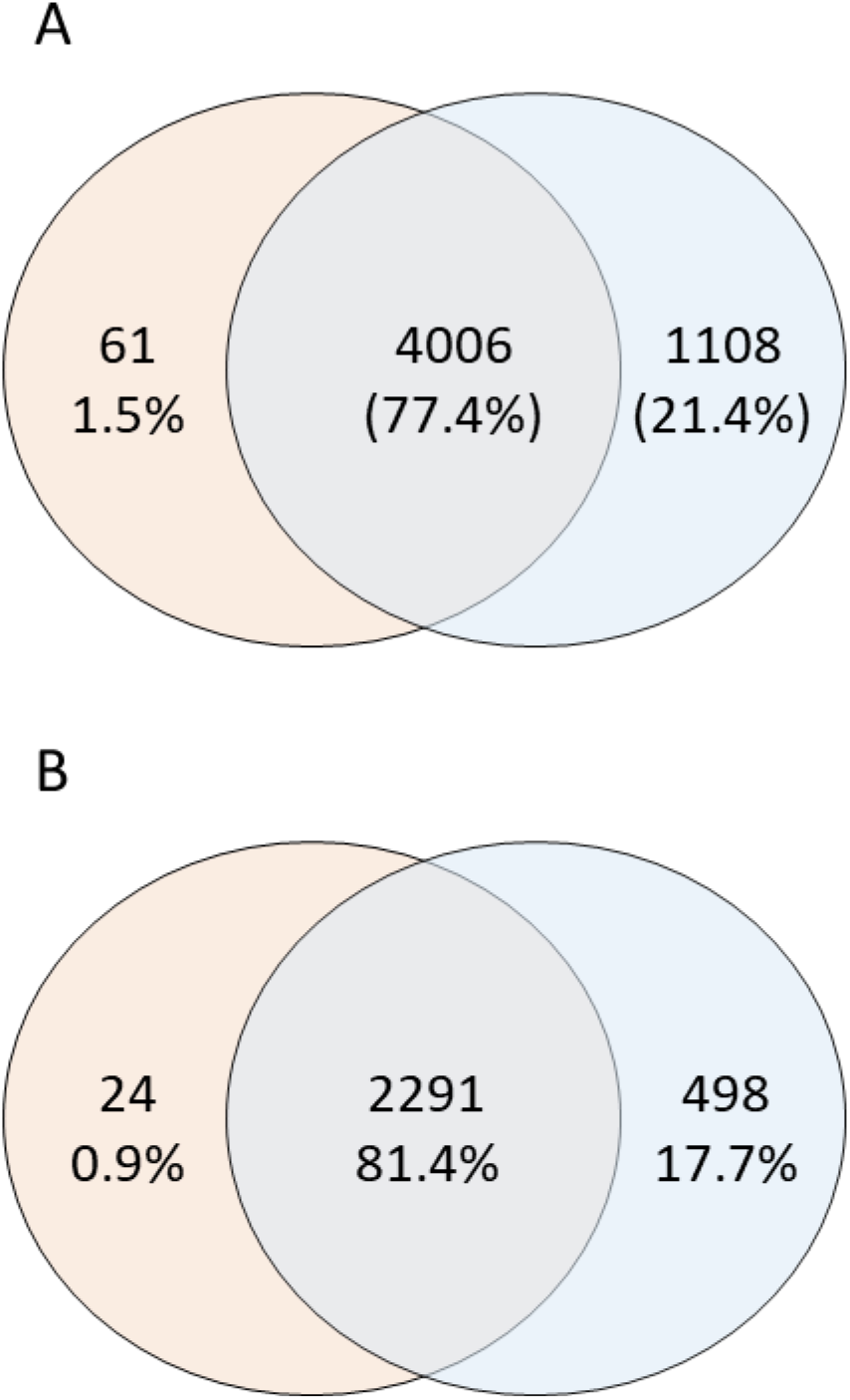
Vann diagram indicating the number of common and sex-specific ESPs (A) and genes related to the SP CpG sites (B) for men (blue) and women (Orange).

### ESP gene enriched for defined biological pathways

To examine the potential links of genes with ESP to aging, we conducted biological pathways enrichment analysis using the Kyoto Encyclopedia of Genes and Genomes (KEGG) pathway database. Fifty-nine pathways were enriched from the ESP gene list with a Benjamini threshold below 0.05 and a fold enrichment of around 2 (Supplementary file 2, Table S4). These pathways were predominantly related to the endocrine system (23%), cancer (13%), neurological communication (10%), cardiovascular system, immune system, cellular focal adhesions and additional pathways that serve multiple biological functions. Table 1 presents a sample of 9 pathways with prominent information regarding their relation to aging. The complete list of age-related pathways which involve genes with ESPs, with the associated ESP genes for each pathway can be found in Supplementary file 2, Table S4.

### ESP Genes known to be associated with aging

We next searched for recognized associations between aging or longevity with ESP genes. We extracted ESP genes from known aging-related genes, which were derived from the GenAge, GenDR, LongevityMap and CellAge databases compiled in The Human Ageing Genomic Resources (HAGR) (33). Of these databases, we identified 149 genes with ESP (for the full list see supplementary file S3, Table S5): 76 genes were from the GenAge, 37 from CellAge, 33 from LongevityMap and 28 from GenDR. A number of genes appeared in more than one database. Hence, 5.3% of all ESP genes were related to aging, not significantly different from the fraction (4.9%) of ageing-related genes derived from these databases (973 genes) out of the total number of genes in the human genome (19,969 genes). While not enriched, then, the list of aging/longevity related genes with ESPs includes a number of genes whose relation to aging is of interest, which are compiled in Table 2. Among the multiple genes with ESPs bearing a particularly significant impact on aging, some, such as ADCY5, APOB, adiponectin, FOXO3, IGF1R, mTORC2, and TXNIP are amongst the most extensively studied in this contest. The overall methylation trend with age, the mean β value and the ESP age of the prominent age-related genes in table 2 are presented for men. The full list of 149 ESP genes linked to aging and longevity with parameters for both genders is provided in tables S5-S7, supplementary file S3. The tables in the supplementary file includes, apart from the parameters presented in table 2, the ESP pattern for each gene and the delta between the first and second calculated slopes (before and after the ESP period).

### Key epigenetic regulators are ESP Genes

Interestingly, we found that key regulators of DNA methylation have ESPs at related CpG sites (Table 3, and supplementary file 3, Table S6), of which the most prominent ones are listed in Table 3. The earliest mean age of appearance of ESP is seen for TRDMT1 and DNMT1 (47, 49 years, respectively), and the latest age of EPS is notable for TET2 (71 years).

**Table 3:**
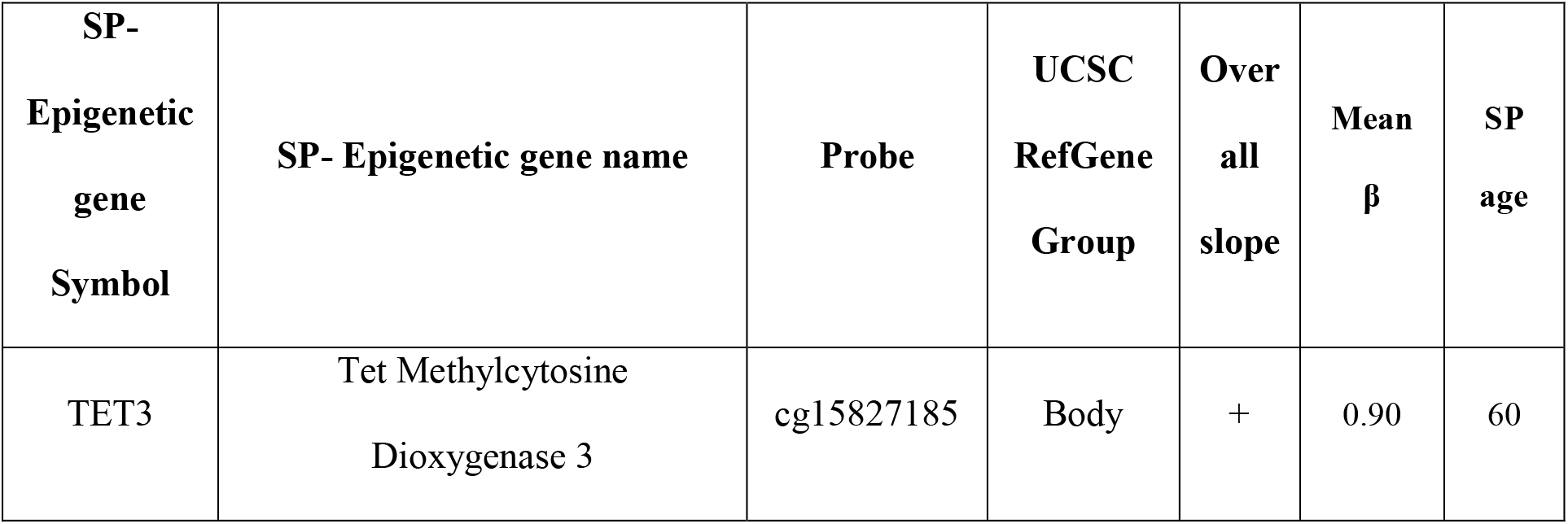
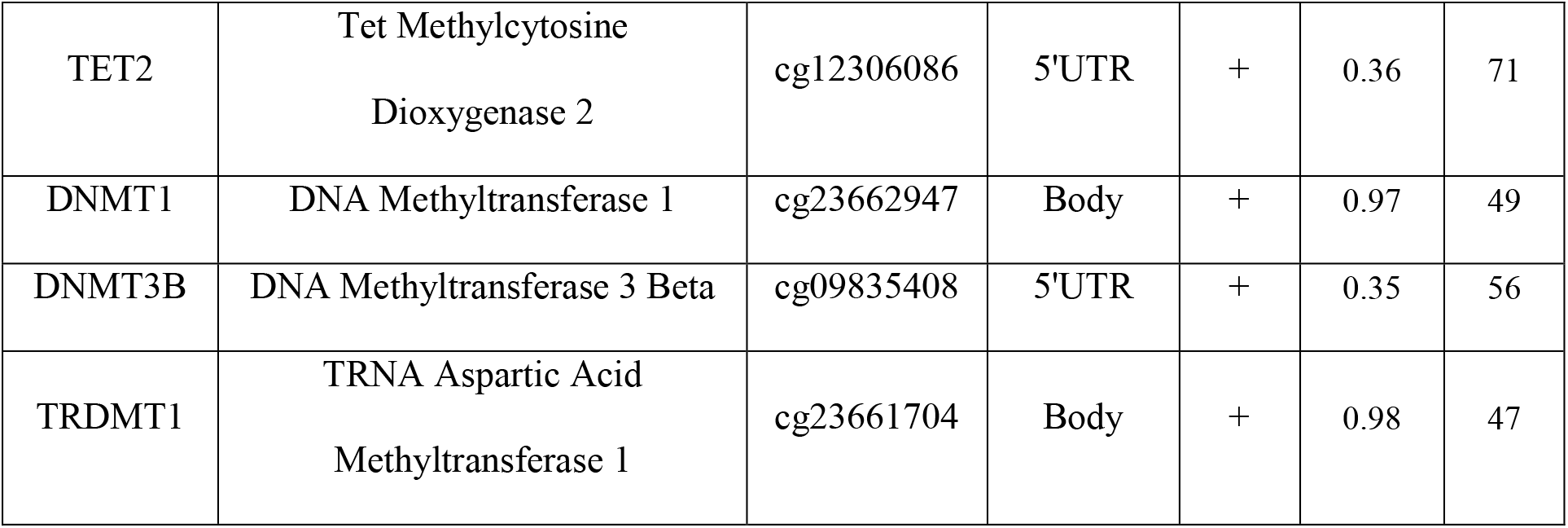
Key SP genes involved in de/methylation processes

| SP-Epigenetic<br>gene<br>Symbol | SP- Epigenetic gene name | Probe | UCSC<br>RefGene<br>Group | Over<br>all<br>slope | Mean<br>$\beta$ | SP<br>age |
| --- | --- | --- | --- | --- | --- | --- |
| TET3 | Tet Methylcytosine<br>Dioxygenase 3 | cg15827185 | Body | + | 0.90 | 60 |
| TET2 | Tet Methylcytosine<br>Dioxygenase 2 | cg12306086 | 5'UTR | + | 0.36 | 71 |
| DNMT1 | DNA Methyltransferase 1 | cg23662947 | Body | + | 0.97 | 49 |
| DNMT3B | DNA Methyltransferase 3 Beta | cg09835408 | 5'UTR | + | 0.35 | 56 |
| TRDMT1 | TRNA Aspartic Acid<br>Methyltransferase 1 | cg23661704 | Body | + | 0.98 | 47 |

### Estrogenic and androgenic SP genes

Because most EPS begin to cluster during the fifth decade of life, in which women experience menopausal transition and hormonal changes begin to afflict men as well, and since the findings in Table 1 show that genes with ESPs are enriched with genes which participate in the estrogen signaling pathway, we queried our ESP gene list regarding the presence and potential enrichment with estrogen and androgen related genes. We found 24 genes that are involved in estrogenic signalling pathway: 14 (of 2813 SP genes) are specifically involved in estrogen receptor alpha (ESR1) signalling. The number of genes related estrogen receptor alpha signalling in the entire 19,969 human gene population, is 55. We calculated an enrichment Chi Square P value of 3.2e-08 and a binomial probability of 0.02 for randomly getting 14 or more genes related to estrogen receptor alpha when taking a sample of 2813 genes. This indicates that there is an enrichment of estrogen receptor alpha-related signalling genes in our ESP list with 1.8fold enrichment. Additionally, we found 28 genes linked to androgen related pathways.

## Discussion

In the present study we set out to assess the relation between age and DNA methylation level in over 2000 subjects collected from 24 different cohorts, whose age spanned from 20 to 80 years. On the average, more than 80% of the CpG sites screened by the Illumina 450K platform which then passed our internal data validation process were found to be stationary over the years. Stationarity does not exclude the possibility of significant intra-individual changes over the years in stationary probes, as the analyzed data is cross-sectional and not based on longitudinal analysis with repeated measurement of methylation of the same probes in the same subjects. Rather, our observations suggest that the majority of the probes included in the 450 k Illumina platform show a stable average methylation fraction between the ages of 20 to 80 years in apparently healthy subjects collected from heterogenous GSE sources.

About one fifth of the validated CpG sites probed in the 450K Illumina platform (69275/341247) were not stationary: they underwent, on the average, consistent changes in methylation level over time that can be described as linear/close to linear, curvilinear or displaying changes in the standard deviation of the mean beta values (methylation fraction). Among the latter group, 5175 non-stationary CpG sites changed linearly or in a manner close to linearity over a certain age span, but then changed course at/around certain age zones, thus displaying a switch point in their trend over the ages of 20-80 years, which we termed an epigenetic switch point (EPS).

In general, switch points detected in the non-stationary probes can be characterized as follows: 1) they are limited to a fraction of the known genes, 2813 of the ∼20,000 human genes. 2) they appear in clusters as three apparent age-related waves: a small wave afflicting ∼100 CpG sites as of the age of 30 to the age of ∼34; a second wave between the ages of 40 to 50 years, composed of∼750 CpG sites which ends 3-4 years earlier in men; and a large wave, including ∼4500CpG sites, between the ages of 50-68 years. 3) their distribution across each of waves grossly follows a normal Gaussian pattern. 4)∼80% of the CpG sites with switch points are shared by women and men; 5) still, there is sexual dimorphism in the distribution of the switch points, such that most of them emerge, peak, decline and vanish earlier in men then in women. Further, in males ∼20% of the ESPs are “male-only” CpGs, compared to only 1.5% “female only” ESPs. Concordantly, switch points occur in less CpG sites and genes in women than in men. Lastly, only men show ESPs with acceleration of an increasing trend in the fraction methylation beyond the ESP-specific age (pattern V, Figure 5).

The clustering of ESPs in defined CpG sites of a limited number of genes, at defined age zones with sexual dimorphism suggests some form of organization rather than a random occurrence. Random switch points would have displayed a spotty pattern spread all over the platform’s probe population, with no predilection for CpG sites, genes, age or sex parameters.

What drives the appearance of EPS clusters in successive waves between the ages of 40-70 years? Could these waves be somehow related to aging-related processes, even though they begin to decline rather sharply as of the late fifties in men and early sixties in women and then entirely fade just a few years later and do not continue to appear in the eight’s decade of life? Certainly, our data cannot provide a fact-based elucidation of this question. However, one cannot escape the consideration of several physiological and clinical changes that precede, succeed or coincide with these waves in contemporary human life: hormonal and metabolic changes and the emergence of cardiovascular morbidities tend to cluster as of the age of 40. In females, menopause does not normally take place until 5 years following the initiation of the second wave, but the menopausal transition is a gradual process lasting 7 up to14 years, most of which take place before the actual cessation of menses (40). Notably, the list of genes with ESP is enriched with ESR1 (estrogen receptor α)-related genes Further, acceleration in metabolic and cardiovascular disease in women actually takes place in post-menopausal women (41,42). Interestingly, ESP in estrogen/estrogen action related genes are nearly all shared by men and women (supplementary file S3, Table S7).

In men, testosterone declines by 0.8% annually as of the age of (43). The decline is much steeper when health status undergoes unfavorable but common impairments (44) with the appearance of overweight, obesity, diabetes and hypertension and cardiovascular events, all of which are on the rise in midlife, particularly after the age of 40 years (40–47). The latter conditions become significant players in females’ health, on the average, about a decade later. Table 1 lists some of the biological pathways that are most strongly enriched with ESPs, in terms of statistical strength (Benjamini’s test with p=0.04-0.00015) and involve processes which are inseparable from the same health trends observed in and as of the fifth decade of life. As examples, we find enrichment with CpG sites with SP in genes linked to adrenergic signaling (hypertension, cardiovascular disease), circadian rhythmicity, cholinergic synapse (obesity, aging, metabolic syndrome), platelet activation (cardiovascular events), insulin secretion (overweight, obesity, diabetes), type II diabetes mellitus and estrogen signaling (menopausal transition).

A close look at the list of genes showing ESPs discloses several examples CpG’s/genes/groups which are of interest vis-à-vis switch points and /or aging. Histone deacetylase 4 (HDAC4) has 7 CpG sites with ESP (Table 2) and interestingly, estrogen act to retain HDAC4 in the nucleus and thus inhibit hypertrophic gene expression and cardiac hypertrophy (48), a common phenotypic phenomenon of heart aging (49). cg15799267, located on the enhancer of arachidonate 15-lipoxygenase B, an enzyme implicated in atherosclerosis (50,51), has an ESP at the age of 45 in women and 54 in men, after which it is increasingly demethylated with age (Table 5S, supplement file 3S).

The presence of ESPs on some of the aging/longevity genes is also intriguing. For example, KO of adenylate cyclase 5 (ADCY5), which involves many G-protein coupled receptor signaling, such as the beta-adrenergic receptor signaling, increased the median life span of mice by 30% (52). Not only was lifespan increased, but resistance to cardiac stress rose whereas age-induced cardiomyopathy and reduction in bone density were lessened (52–54). The co-presence of ESPs in the EGFR (epidermal factor receptor), insulin receptor, IGF1R, IG2R, is also notable, as these are network players at the interface of growth control (including cancer growth) and vascular aging with metabolic carbohydrate, fat and protein homeostasis, where both excess activity or improper activation can modulate metabolic disease, bridge insulin resistance to cancer and thus affect survival (55–58). Disruption of the insulin/insulin-like growth factor 1 signaling (IIS) pathway in Caenorhabditis elegans was found to double its lifespan (59). At the organ level, IGF1R KO in mice cardiomyocytes attenuates cardiac hypertrophy associated with cardiac aging (60) whereas constitutive activation of IGF1R in vascular smooth muscle cells of old mice accelerates the development of atherosclerosis (61). <u>The IR/IGF-1</u> signaling cascade also negatively regulates Forkhead Box O3 (FOXO3), a transcription factor with a strong positive impact on aging and age-related phenotypes, operating through enhancement of cellular ability to sustain stress (62). Concordant with this role of FOXO3, natural SNPs in the FOXO3 gene were associated with increased longevity of American men of Japanese ancestry aged ≥95 years (61,62).

The Mechanistic Target of Rapamycin Kinase (mTOR) is tightly linked to aging biology (65–68). It functions as an intracellular energy sensor and a central regulator of growth, proliferation, metabolism, survival, protein synthesis, apoptosis autophagy and transcription (69). Impressively, rapamycin, an inhibitor of the mTOR complex 1 (mTORC1) has thus far increased lifespan in all model organisms studied (65). The protein complex mTOR2 promotes the activation of insulin receptors and insulin-like growth factor 1 receptors and as such, disruption of mTORC2 leads to glucose intolerance, diabetes, lower activity level and immunosuppression (66,70). Its activity is inhibited by insulin/insulin-like growth factor (IGF-1) signaling (IIS) cascade stimulation. Finally, differential methylation in CpG sites related to Thioredoxin Interacting Protein (TXNIP), shown here to have ESPs, were found in Type 2 Diabetes population, commonly, associated with aging (71–73).

Table 3 highlights 5 ESPs on genes encoding enzymes which directly affect DNA methylation. DNMT1, DNMT3B are classical writer enzymes responsible for DNA methylations. DNMT1, catalyzes the transfer of a methyl group to a cytosine nucleotide and is mainly responsible for maintaining DNA methylation, which ensures the fidelity of this epigenetic patterns across cell divisions. In contrast, DNA Methyltransferase 3 Beta (DNMT3B) functions in *de novo* methylation. Hence, a change at the ESP for DNMT3B at a mean age of 56 may be consequential. TRDMT1, tRNA aspartic acid methyltransferase 1, also known as DNMT2, can also act as a writer, but is importantly linked to resistance to stress, including oxidative stress, inflammation, salt stress, and cellular senescence (74,75). TET enzymes generally function as 5-mC erasers by the conversion of 5-mC into 5-hmC, an intermediary metabolite that releases the presumed effect of the methylated CpG site. Notably, 2 of the three known TET enzymes, TET3 and TET2 are shown to possess ESP age of 71 and 60 years, respectively. Hence, some ESPs can be identified in the DNA methylation/demethylation machinery itself, preceding or coinciding with the overall ESPs waves)Table 3).

Because these are just few of the >2800 genes with SP, and since ESPs waves encompass a distinct, but broad age ranges, a detailed analysis of the of potential specific potential links between the unfolding clinical unfolding of physiological processes and common diseases in evolution and the epigenetic waves and their individual components comprises a formidable challenge for future work.

At the present phase, the conceptualization of the ESPs waves as a potentially significant phenomenon, requires consideration of the fact that whereas aging itself, including epigenetic aging (5–7,14) goes on continuously with advancing years, the ESPs waves apparently begin to wane after the age of ∼58 in men and ∼63 in women and then completely disappear within the following 5-8 years. This transient pattern of epigenetic waves might indicate that the CpG signals comprising the ESP waves are mechanistically involved in or at the least echo some biological resetting that permits aging to advance, but may also reflect compensatory pro-longevity biosystems activated by signals of aging and senescence. It is also possible, particularly since we analyze age by cross-sectional and not longitudinal tools, that the waning of the ESPs reflects a Darwinian selection, the survival of the fittest: subjects who remained alive to their 8th decade may have not experienced the switch point waves that afflicted non-survivors, thus leaving the 8th decade of life free of ESP waves. While potentially testable, these hypotheses cannot be addressed without in depth longitudinal studies.

In conclusion, this is the first report of epigenetic aging waves in the form of clustering of EPS in a fraction of the genes comprising the human genome, in an age and sex specific manner. The main weakness of this report is that it is based on cross sectional analysis of mean methylation level and not on longitudinal observations in the same individuals and therefore does not deal with intra-individual changes in DNA methylation. This novel phenomenon, however, which appears well organized and is linked to several transitional periods in human life, lays the foundations to future quests to understand its underlying triggers as well as health and aging-related consequences.

## List of abbreviations

ESP: epigenetic switch points
HDAC4: Histone deacetylase 4
mTOR: Mechanistic Target of Rapamycin Kinase
FOXO3: Forkhead Box O3
TXNIP: Thioredoxin Interacting Protein
ADCY5: adenylate cyclase 5
IGFR: insulin growth factor receptor
IGF: insulin growth factor
IIS: insulin/insulin-like growth factor (IGF-1) signaling
DNMT1: DNA methyl transferase 1
TRDMT1: tRNA aspartic acid methyltransferase 1, also known as DNMT2,
DNMT3B: Methyltransferase 3 Beta
TET1/2: Ten-Eleven Translocation 1/2

## Ethics approval and consent to participate

Not applicable

## Consent for publication

Not applicable

## Availability of data and materials

The GEO number of the data sets analyzed during the current study are listed in Additional file 1, table S1.

## Author contribution statement

N.S. initiated the project. N.S., E.S., T.K and T.S. designed the study. T.K, E.S, T.S, Y.M, and G.S have selected and filtered the data source. T.K and E.S did all the mathematical and statistical calculations. Y.E and M.P gave useful reviews and comments on the paper. N.S, E.S and T.S. wrote the manuscript. All authors gave approval to the final version of the manuscript.

## Acknowledgment

We would like to thank Sami and Tova Sagol for their generous donation to the Sagol Epigenetic center, which made this study possible.

## Supplementary material

Supplementary file 1: table S1: The GEO number of the data sets analyzed during the current study, table S2: the 19 parameters used in the decision tree model, figure S1: Age distribution of the sample dataset by gender, figure S2: Distribution of β values in type III ESPs.

Supplementary file 2: Table 3: The complete list of 5175 ESP, Table 4: The complete list of age-related pathways which involve genes with ESPs.

Supplementary file 3: Table 5: The complete list of 149 ESP genes which are aging/ longevity genes. Table 6: DNA methylation process-related ESP. Table 7: Estrogen related ESP. Table 8: Androgen related ESP.

## References

1. Mays-Hoopes LL. DNA Methylation in Aging and Cancer. J Gerontol [Internet]. 1989 Nov 1 [cited 2022 Sep 29];44(6):35–6. Available from: https://academic.oup.com/geronj/article/44/6/35/564452

2. Jones MJ, Goodman SJ, Kobor MS. DNA methylation and healthy human aging.2015 Dec 1 [cited 2021 Oct 4];14(6):924–32. Available from: https://pubmed.ncbi.nlm.nih.gov/25913071/

3. Drinkwater RD, Blake TJ, Morley AA, Turner DR. Human lymphocytes aged in vivo have reduced levels of methylation in transcriptionally active and inactive DNA. Mutat Res [Internet]. 1989 [cited 2022 Sep 30];219(1):29–37. Available from: https://pubmed.ncbi.nlm.nih.gov/2911269/

4. Horvath S. DNA methylation age of human tissues and cell types. Genome Biol. 2013 Oct 21;14(10):R115.

5. Horvath S, Zhang Y, Langfelder P, Kahn RS, Boks MP, Eijk K van, et al. Aging effects on DNA methylation modules in human brain and blood tissue. Genome Biol [Internet]. 2012 [cited 2022 Sep 30];13(10):R97. Available from: https://www.academia.edu/11469305/Aging_effects_on_DNA_methylation_modules_in_human_brain_and_blood_tissue

6. Johansson Å, Enroth S, Gyllensten U. Continuous Aging of the Human DNA Methylome Throughout the Human Lifespan. PLoS One [Internet]. 2013 Jun 27 [cited 2022 Sep 29];8(6):e67378. Available from: https://journals.plos.org/plosone/article?id=10.1371/journal.pone.0067378

7. Florath I, Butterbach K, Müller H, Bewerunge-hudler M, Brenner H. Cross-sectional and longitudinal changes in DNA methylation with age: an epigenome-wide analysis revealing over 60 novel age-associated CpG sites. Hum Mol Genet [Internet]. 2014 Mar [cited 2022 Sep 29];23(5):1186–201. Available from: https://pubmed.ncbi.nlm.nih.gov/24163245/

8. Illingworth RS, Bird AP. CpG islands--’a rough guide’. FEBS Lett [Internet]. 2009 Jun 5 [cited 2022 Sep 29];583(11):1713–20. Available from: https://pubmed.ncbi.nlm.nih.gov/19376112/

9. Teschendorff AE, West J, Beck S. Age-associated epigenetic drift: implications, and a case of epigenetic thrift? Hum Mol Genet [Internet]. 2013 Oct [cited 2022 Sep 29];22(R1). Available from: https://pubmed.ncbi.nlm.nih.gov/23918660/

10. Saxonov S, Berg P, Brutlag DL. A genome-wide analysis of CpG dinucleotides in the human genome distinguishes two distinct classes of promoters. Proc Natl Acad Sci U S A [Internet]. 2006 Jan 31 [cited 2022 Sep 29];103(5):1412–7. Available from: https://www.pnas.org/doi/abs/10.1073/pnas.0510310103

11. Hannum G, Guinney J, Zhao L, Zhang L, Hughes G, Sadda SV, et al. Genome-wide Methylation Profiles Reveal Quantitative Views of Human Aging Rates. Mol Cell [Internet]. 2013 Jan 24 [cited 2021 Oct 4];49(2):359–67. Available from: https://pubmed.ncbi.nlm.nih.gov/23177740/

12. Weidner CI, Lin Q, Koch CM, Eisele L, Beier F, Ziegler P, et al. Aging of blood can be tracked by DNA methylation changes at just three CpG sites. Genome Biol [Internet]. 2014 Feb 3 [cited 2022 Sep 29];15(2):1–12. Available from: https://genomebiology.biomedcentral.com/articles/10.1186/gb-2014-15-2-r24

13. Mcclay JL, Aberg KA, Clark SL, Nerella S, Kumar G, Xie LY, et al. A methylome-wide study of aging using massively parallel sequencing of the methyl-CpG-enriched genomic fraction from blood in over 700 subjects. Hum Mol Genet [Internet]. 2014 Mar [cited 2022 Sep 29];23(5):1175–85. Available from: https://pubmed.ncbi.nlm.nih.gov/24135035/

14. Shahal T, Segev E, Konstantinovsky T, Marcus Y, Shefer G, Pasmanik-Chor M, et al. Deconvolution of the epigenetic age discloses distinct inter-personal variability in epigenetic aging patterns. Epigenetics and Chromatin [Internet]. 2022 Dec 1 [cited 2022 Sep 29];15(1):1–12. Available from: https://epigeneticsandchromatin.biomedcentral.com/articles/10.1186/s13072-022-00441-y

15. Chen BH, Marioni RE, Colicino E, Peters MJ, Ward-Caviness CK, Tsai PC, et al. DNA methylation-based measures of biological age: Meta-analysis predicting time to death. Aging (Albany NY) [Internet]. 2016 [cited 2022 Sep 29];8(9):1844–65. Available from: https://pubmed.ncbi.nlm.nih.gov/27690265/

16. Horvath S, Gurven M, Levine ME, Trumble BC, Kaplan H, Allayee H, et al. An epigenetic clock analysis of race/ethnicity, sex, and coronary heart disease. Genome Biol [Internet]. 2016 Aug 11 [cited 2021 Oct 4];17(1):171. Available from: https://genomebiology.biomedcentral.com/articles/10.1186/s13059-016-1030-0

17. Zhang Y, Wilson R, Heiss J, Breitling LP, Saum KU, Schöttker B, et al. DNA methylation signatures in peripheral blood strongly predict all-cause mortality. Nat Commun. 2017;8:14617.

18. Jain P, Binder AM, Chen B, Parada H, Gallo LC, Alcaraz J, et al. Analysis of Epigenetic Age Acceleration and Healthy Longevity Among Older US Women. JAMA Netw Open [Internet]. 2022 Jul 1 [cited 2022 Sep 29];5(7):e2223285–e2223285. Available from: https://jamanetwork.com/journals/jamanetworkopen/fullarticle/2794706

19. Levine ME, Hosgood HD, Chen B, Absher D, Assimes T, Horvath S. DNA methylation age of blood predicts future onset of lung cancer in the women’s health initiative. Aging (Albany NY) [Internet]. 2015 [cited 2022 Sep 29];7(9):690–700. Available from: https://pubmed.ncbi.nlm.nih.gov/26411804/

20. Ambatipudi S, Horvath S, Perrier F, Cuenin C, Hernandez-Vargas H, Le Calvez-Kelm F, et al. DNA methylome analysis identifies accelerated epigenetic ageing associated with postmenopausal breast cancer susceptibility. Eur J Cancer [Internet]. 2017 Apr 1 [cited 2022 Sep 29];75:299–307. Available from: https://pubmed.ncbi.nlm.nih.gov/28259012/

21. Roetker NS, Pankow JS, Bressler J, Morrison AC, Boerwinkle E. Prospective Study of Epigenetic Age Acceleration and Incidence of Cardiovascular Disease Outcomes in the ARIC Study (Atherosclerosis Risk in Communities). Circ Genomic Precis Med [Internet]. 2018 Mar 1 [cited 2022 Sep 29];11(3):e001937. Available from: https://pubmed.ncbi.nlm.nih.gov/29555670/

22. Snir S, Farrell C, Pellegrini M. Human epigenetic ageing is logarithmic with time across the entire lifespan. Epigenetics [Internet]. 2019 Sep 2 [cited 2022 Sep 30];14(9):912–26. Available from: https://pubmed.ncbi.nlm.nih.gov/31138013/

23. Rubbi L, Zhang H, Feng J, He C, Kurnia P, Ratan P, et al. The effects of age, sex, weight, and breed on canid methylomes. Epigenetics [Internet]. 2022 [cited 2022 Sep 30]; Available from: https://pubmed.ncbi.nlm.nih.gov/35502722/

24. Martino DJ, Tulic MK, Gordon L, Hodder M, Richman T, Metcalfe J, et al. Evidence for age-related and individual-specific changes in DNA methylation profile of mononuclear cells during early immune development in humans. Epigenetics [Internet]. 2011 [cited 2022 Sep 29];6(9):1085–94. Available from: https://pubmed.ncbi.nlm.nih.gov/21814035/

25. Herbstman JB, Wang S, Perera FP, Lederman SA, Vishnevetsky J, Rundle AG, et al. Predictors and consequences of global DNA methylation in cord blood and at three years. PLoS One [Internet]. 2013 Sep 4 [cited 2022 Sep 29];8(9). Available from: https://pubmed.ncbi.nlm.nih.gov/24023780/

26. Han L, Zhang H, Kaushal A, Rezwan FI, Kadalayil L, Karmaus W, et al. Changes in DNA methylation from pre-to post-adolescence are associated with pubertal exposures. Clin Epigenetics [Internet]. 2019 Dec 2 [cited 2022 Sep 29];11(1):1–14. Available from: https://clinicalepigeneticsjournal.biomedcentral.com/articles/10.1186/s13148-019-0780-4

27. Morris TJ, Butcher LM, Feber A, Teschendorff AE, Chakravarthy AR, Wojdacz TK, et al. ChAMP: 450k Chip Analysis Methylation Pipeline. Bioinformatics [Internet]. 2014 Feb 1 [cited 2022 Sep 29];30(3):428–30. Available from: https://academic.oup.com/bioinformatics/article/30/3/428/228299

28. Teschendorff AE, Marabita F, Lechner M, Bartlett T, Tegner J, Gomez-Cabrero D, et al. A beta-mixture quantile normalization method for correcting probe design bias in Illumina Infinium 450 k DNA methylation data. 2013 Jan 15 [cited 2022 Sep 29];29(2):189–96. Available from: https://academic.oup.com/bioinformatics/article/29/2/189/204142

29. Houseman EA, Accomando WP, Koestler DC, Christensen BC, Marsit CJ, Nelson HH, et al. DNA methylation arrays as surrogate measures of cell mixture distribution. BMC Bioinformatics [Internet]. 2012 May 8 [cited 2022 Sep 30];13(1):1–16. Available from: https://bmcbioinformatics.biomedcentral.com/articles/10.1186/1471-2105-13-86

30. Robert B. Cleveland, William S. Cleveland, Jean E. McRae IT. STL: A Seasonal-Trend Decomposition Procedure Based on Loess - ProQuest [Internet]. Journal of official statistics. 1990 [cited 2022 Sep 30]. p. 3–73. Available from: https://www.proquest.com/docview/1266805989?parentSessionId=KCOre8G8D2YY70EegFfTWWxf7R6R6ipCvz%2Fezt%2Fxa%2BY%3D

31. Kanehisa M, Goto S, Sato Y, Furumichi M, Tanabe M. KEGG for integration and interpretation of large-scale molecular data sets. Nucleic Acids Res [Internet]. 2012 Jan [cited 2022 Sep 30];40(Database issue). Available from: https://pubmed.ncbi.nlm.nih.gov/22080510/

32. Lynch DR, Farmer JM, Balcer LJ, Wilson RB. Friedreich ataxia: Effects of genetic understanding on clinical evaluation and therapy. Vol. 59, Archives of Neurology. 2002. p. 743–7.

33. Tacutu R, Thornton D, Johnson E, Budovsky A, Barardo D, Craig T, et al. Human Ageing Genomic Resources: new and updated databases. Jan [cited 2022 Sep 20];46(D1). Available from: https://pubmed.ncbi.nlm.nih.gov/29121237/

34. Fernandes M, Wan C, Tacutu R, Barardo D, Rajput A, Wang J, et al. Systematic analysis of the gerontome reveals links between aging and age-related diseases. 2016 Nov 1 [cited 2022 Sep 29];25(21):4804–18. Available from: https://pubmed.ncbi.nlm.nih.gov/28175300/

35. De Magalhães JP, Toussaint O. GenAge: A genomic and proteomic network map of human ageing. FEBS Lett [Internet]. 2004 Jul 30 [cited 2022 Sep 30];571(1–3):243–7. Available from: https://pubmed.ncbi.nlm.nih.gov/15280050/

36. Plank M, Wuttke D, Van Dam S, Clarke SA, D. Magalhães JP. A meta-analysis of caloric restriction gene expression profiles to infer common signatures and regulatory mechanisms. Mol Biosyst [Internet]. 2012 Mar [cited 2022 Sep 29];8(4):1339–49. Available from: https://pubmed.ncbi.nlm.nih.gov/22327899/

37. Wuttke D, Connor R, Vora C, Craig T, Li Y, Wood S, et al. Dissecting the gene network of dietary restriction to identify evolutionarily conserved pathways and new functional genes. PLoS Genet [Internet]. 2012 Aug [cited 2022 Sep 29];8(8). Available from: https://pubmed.ncbi.nlm.nih.gov/22912585/

38. Avelar RA, Ortega JG, Tacutu R, Tyler EJ, Bennett D, Binetti P, et al. A multidimensional systems biology analysis of cellular senescence in aging and disease. Genome Biol [Internet]. 2020 Apr 7 [cited 2022 Sep 30];21(1). Available from: https://pubmed.ncbi.nlm.nih.gov/32264951/

39. Oliveros, J.C. (2007-2015) Venny. An Interactive Tool for Comparing Lists with Venn’s Diagrams. https://bioinfogp.cnb.csic.es/tools/venny/index.html.

40. El Khoudary SR, Greendale G, Crawford SL, Avis NE, Brooks MM, Thurston RC, et al. The menopause transition and women’s health at midlife: a progress report from the Study of Women’s Health Across the Nation (SWAN). Menopause [Internet]. 2019 Oct 1 [cited 2022 Sep 30];26(10):1213. Available from: /pmc/articles/PMC6784846/

41. Miller JJ, Heather LC. Cardiometabolic risk factors vary with age differently in females and males. Nat Cardiovasc Res 2022 19 [Internet]. 2022 Sep 12 [cited 2022 Sep 30];1(9):796–7. Available from: https://www.nature.com/articles/s44161-022-00130-9

42. Gerdts E, Regitz-Zagrosek V. Sex differences in cardiometabolic disorders. Nat Med 2019 2511 [Internet]. 2019 Nov 7 [cited 2022 Sep 29];25(11):1657–66. Available from: https://www.nature.com/articles/s41591-019-0643-8

43. Feldman HA, Longcope C, Derby CA, Johannes CB, Araujo AB, Coviello AD, et al. Age trends in the level of serum testosterone and other hormones in middle-aged men: longitudinal results from the Massachusetts male aging study. J Clin Endocrinol Metab [Internet]. 2002 [cited 2022 Sep 29];87(2):589–98. Available from: https://pubmed.ncbi.nlm.nih.gov/11836290/

44. Hildrum B, Mykletun A, Hole T, Midthjell K, Dahl AA. Age-specific prevalence of the metabolic syndrome defined by the International Diabetes Federation and the National Cholesterol Education Program: The Norwegian HUNT 2 study. BMC Public Health [Internet]. 2007 Aug 29 [cited 2022 Sep 29];7(1):1–9. Available from: https://bmcpublichealth.biomedcentral.com/articles/10.1186/1471-2458-7-220

45. Saeedi P, Petersohn I, Salpea P, Malanda B, Karuranga S, Unwin N, et al. Global and regional diabetes prevalence estimates for 2019 and projections for 2030 and 2045: Results from the International Diabetes Federation Diabetes Atlas, 9th edition. Diabetes Res Clin Pract [Internet]. 2019 Nov 1 [cited 2022 Sep 29];157. Available from: https://pubmed.ncbi.nlm.nih.gov/31518657/

46. Meyer MR, Haas E, Barton M. Gender Differences of Cardiovascular Disease. Hypertension [Internet]. 2006 Jun 1 [cited 2022 Sep 30];47(6):1019–26. Available from: https://www.ahajournals.org/doi/abs/10.1161/01.HYP.0000223064.62762.0b

47. Tsay YC, Chen CH, Pan WH. Ages at Onset of 5 Cardiometabolic Diseases Adjusting for Nonsusceptibility: Implications for the Pathogenesis of Metabolic Syndrome. Am J Epidemiol [Internet]. 2016 Sep 1 [cited 2022 Sep 29];184(5):366–77. Available from: https://academic.oup.com/aje/article/184/5/366/2388960

48. Pedrama A, Razandi M, Narayanan R, Dalton JT, McKinsey TA, Levin ER. Estrogen regulates histone deacetylases to prevent cardiac hypertrophy. Mol Biol Cell [Internet]. 2013 Dec 15 [cited 2022 Sep 29];24(24):3805–18. Available from: https://www.molbiolcell.org/doi/10.1091/mbc.e13-08-0444

49. Sessions AO, Engler AJ. Mechanical Regulation of Cardiac Aging in Model Systems. Circ Res [Internet]. 2016 May 13 [cited 2022 Sep 29];118(10):1553–62. Available from: https://pubmed.ncbi.nlm.nih.gov/27174949/

50. Vijil C, Hermansson C, Jeppsson A, Bergström G, Hultén LM. Arachidonate 15-Lipoxygenase Enzyme Products Increase Platelet Aggregation and Thrombin Generation. PLoS One [Internet]. 2014 Feb 12 [cited 2022 Sep 29];9(2):e88546. Available from: https://journals.plos.org/plosone/article?id=10.1371/journal.pone.0088546

51. Gertow K, Nobili E, Folkersen L, Newman JW, Pedersen TL, Ekstrand J, et al. 12- and 15-lipoxygenases in human carotid atherosclerotic lesions: Associations with cerebrovascular symptoms. Atherosclerosis. 2011 Apr 1;215(2):411–6.

52. Yan L, Vatner DE, O’Connor JP, Ivessa A, Ge H, Chen W, et al. Type 5 adenylyl cyclase disruption increases longevity and protects against stress. Cell [Internet]. 2007 Jul 27 [cited 2022 Sep 30];130(2):247–58. Available from: https://pubmed.ncbi.nlm.nih.gov/17662940/

53. Vatner SF, Yan L, Ishikawa Y, Vatner DE, Sadoshima J. Adenylyl cyclase type 5 disruption prolongs longevity and protects the heart against stress. Circ J [Internet]. 2009 [cited 2022 Sep 29];73(2):195–200. Available from: https://pubmed.ncbi.nlm.nih.gov/19106458/

54. Vatner SF, Pachon RE, Vatner DE. Inhibition of adenylyl cyclase type 5 increases longevity and healthful aging through oxidative stress protection. Oxid Med Cell Longev [Internet]. 2015 [cited 2022 Sep 29];2015. Available from: https://pubmed.ncbi.nlm.nih.gov/25945149/

55. Yakar S, Adamo ML. Insulin-like growth factor 1 physiology: lessons from mouse models. Endocrinol Metab Clin North Am [Internet]. 2012 Jun [cited 2022 Sep 30];41(2):231–47. Available from: https://pubmed.ncbi.nlm.nih.gov/22682628/

56. LeRoith D, Yakar S. Mechanisms of Disease: metabolic effects of growth hormone and insulin-like growth factor 1. Nat Clin Pract Endocrinol Metab 2007 33 [Internet]. 2007 Mar [cited 2022 Sep 30];3(3):302–10. Available from: https://www.nature.com/articles/ncpendmet0427

57. Rosenfeld RG. Insulin-like Growth Factors and the Basis of Growth. https://doi.org/101056/NEJMp038156 [Internet]. 2003 Dec 4 [cited 2022 Sep 30];349(23):2184–6. Available from: https://www.nejm.org/doi/full/10.1056/NEJMp038156

58. Pawlikowska L, Hu D, Huntsman S, Sung A, Chu C, Chen J, et al. Association of common genetic variation in the insulin/IGF1 signaling pathway with human longevity. Aging Cell [Internet]. 2009 [cited 2022 Sep 30];8(4):460–72. Available from: https://pubmed.ncbi.nlm.nih.gov/19489743/

59. Kenyon C, Chang J, Gensch E, Rudner A, Tabtiang R. A C. elegans mutant that lives twice as long as wild type. Nature [Internet]. 1993 [cited 2022 Sep 30];366(6454):461–4. Available from: https://pubmed.ncbi.nlm.nih.gov/8247153/

60. Ock S, Lee WS, Ahn J, Kim HM, Kang H, Kim HS, et al. Deletion of IGF-1 Receptors in Cardiomyocytes Attenuates Cardiac Aging in Male Mice. Endocrinology [Internet]. 2016 Jan 1 [cited 2022 Sep 30];157(1):336–45. Available from: https://pubmed.ncbi.nlm.nih.gov/26469138/

61. Li M, Chiu JF, Gagne J, Fukagawa NK. Age-related differences in insulin-like growth factor-1 receptor signaling regulates Akt/FOXO3a and ERK/Fos pathways in vascular smooth muscle cells. J Cell Physiol [Internet]. 2008 Nov 1 [cited 2022 Sep 30];217(2):377–87. Available from: https://onlinelibrary.wiley.com/doi/full/10.1002/jcp.21507

62. Calnan DR, Brunet A. The FoxO code. Oncogene [Internet]. 2008 Apr 7 [cited 2022 Sep 30];27(16):2276–88. Available from: https://pubmed.ncbi.nlm.nih.gov/18391970/

63. Willcox BJ, Donlon TA, He Q, Chen R, Grove JS, Yano K, et al. FOXO3A genotype is strongly associated with human longevity. Proc Natl Acad Sci U S A [Internet]. 2008 Sep 16 [cited 2022 Sep 30];105(37):13987–92. Available from: https://www.pnas.org/doi/abs/10.1073/pnas.0801030105

64. Flachsbart F, Caliebe A, Kleindorp R, Blanché H, Von Eller-Eberstein H, Nikolaus S, et al. Association of FOXO3A variation with human longevity confirmed in German centenarians. Proc Natl Acad Sci U S A [Internet]. 2009 Feb 2 [cited 2022 Sep 30];106(8):2700. Available from: /pmc/articles/PMC2650329/

65. Weichhart T. mTOR as Regulator of Lifespan, Aging, and Cellular Senescence: A Mini-Review. Gerontology [Internet]. 2018 Feb 1 [cited 2022 Sep 29];64(2):127–34. Available from: https://pubmed.ncbi.nlm.nih.gov/29190625/

66. Arriola Apelo SI, Lamming DW. Rapamycin: An InhibiTOR of Aging Emerges From the Soil of Easter Island. J Gerontol A Biol Sci Med Sci [Internet]. 2016 Jul 1 [cited 2022 Sep 29];71(7):841–9. Available from: https://pubmed.ncbi.nlm.nih.gov/27208895/

67. Papadopoli D, Boulay K, Kazak L, Pollak M, Mallette FA, Topisirovic I, et al. mTOR as a central regulator of lifespan and aging. F1000Research [Internet]. 2019 [cited 2022 Sep 29];8. Available from: /pmc/articles/PMC6611156/

68. Yu M, Zhang H, Wang B, Zhang Y, Zheng X, Shao B, et al. Key Signaling Pathways in Aging and Potential Interventions for Healthy Aging. Cells [Internet]. 2021 Mar 1 [cited 2022 Sep 29];10(3):1–26. Available from: /pmc/articles/PMC8002281/

69. Albert V, Hall MN. mTOR signaling in cellular and organismal energetics. Curr Opin Cell Biol [Internet]. 2015 [cited 2022 Sep 29];33(1):55–66. Available from: https://pubmed.ncbi.nlm.nih.gov/25554914/

70. Chellappa K, Brinkman JA, Mukherjee S, Morrison M, Alotaibi MI, Carbajal KA, et al. Hypothalamic mTORC2 is essential for metabolic health and longevity. Aging Cell [Internet]. 2019 Oct 1 [cited 2022 Sep 29];18(5). Available from: /pmc/articles/PMC6718533/

71. Juvinao-Quintero DL, Marioni RE, Ochoa-Rosales C, Russ TC, Deary IJ, van Meurs JBJ, et al. DNA methylation of blood cells is associated with prevalent type 2 diabetes in a meta-analysis of four European cohorts. Clin Epigenetics [Internet]. 2021 Dec 1 [cited 2022 Sep 30];13(1):1–14. Available from: https://clinicalepigeneticsjournal.biomedcentral.com/articles/10.1186/s13148-021-01027-3

72. Cardona A, Day FR, Perry JRB, Loh M, Chu AY, Lehne B, et al. Epigenome-wide association study of incident type 2 diabetes in a British population: EPIC-Norfolk study. Diabetes [Internet]. 2019 Dec 1 [cited 2022 Sep 30];68(12):2315–26. Available from: https://pubmed.ncbi.nlm.nih.gov/31506343/

73. Walaszczyk E, Luijten M, Spijkerman AMW, Bonder MJ, Lutgers HL, Snieder H, et al. DNA methylation markers associated with type 2 diabetes, fasting glucose and HbA1c levels: a systematic review and replication in a case-control sample of the Lifelines study. Diabetologia [Internet]. 2018 Feb 1 [cited 2022 Sep 30];61(2):354–68. Available from: https://pubmed.ncbi.nlm.nih.gov/29164275/

74. Lewinska A, Adamczyk-Grochala J, Kwasniewicz E, Wnuk M. Downregulation of methyltransferase Dnmt2 results in condition-dependent telomere shortening and senescence or apoptosis in mouse fibroblasts. J Cell Physiol [Internet]. 2017 Dec 1 [cited 2022 Sep 29];232(12):3714–26. Available from: https://pubmed.ncbi.nlm.nih.gov/28177119/

75. Li Z, Qi X, Zhang X, Yu L, Gao L, Kong W, et al. TRDMT1 exhibited protective effects against LPS-induced inflammation in rats through TLR4-NF-κB/MAPK-TNF-α pathway. Anim Model Exp Med [Internet]. 2022 Apr 1 [cited 2022 Sep 29];5(2):172–82. Available from: https://pubmed.ncbi.nlm.nih.gov/35474613/

