## Supplementary file S1 for "Epigenetic aging waves: Artificial intelligence detects clustering of switch points in DNA methylation rate in defined sex-dependent age periods"

Table S1: data sets GSE code and number of samples from each data set used for farther methylation ESP analysis

| **GSE** | **Number of Samples** |
| --- | --- |
| GSE87571 | 572 |
| GSE72775 | 221 |
| GSE111629 | 197 |
| GSE69270 | 166 |
| GSE50660 | 163 |
| GSE106648 | 131 |
| GSE128235 | 118 |
| GSE74414 | 95 |
| GSE77445 | 81 |
| GSE64495 | 73 |
| GSE67751 | 53 |
| GSE87648 | 52 |
| GSE113725 | 48 |
| GSE52588 | 41 |
| GSE41169 | 34 |
| GSE77056 | 24 |
| GSE52114 | 20 |
| GSE85506 | 18 |
| GSE107737 | 11 |
| GSE53130 | 9 |
| GSE107143 | 8 |
| GSE99624 | 5 |
| GSE84003 | 2 |
| GSE111165 | 1 |
| **Total** | **2143** |

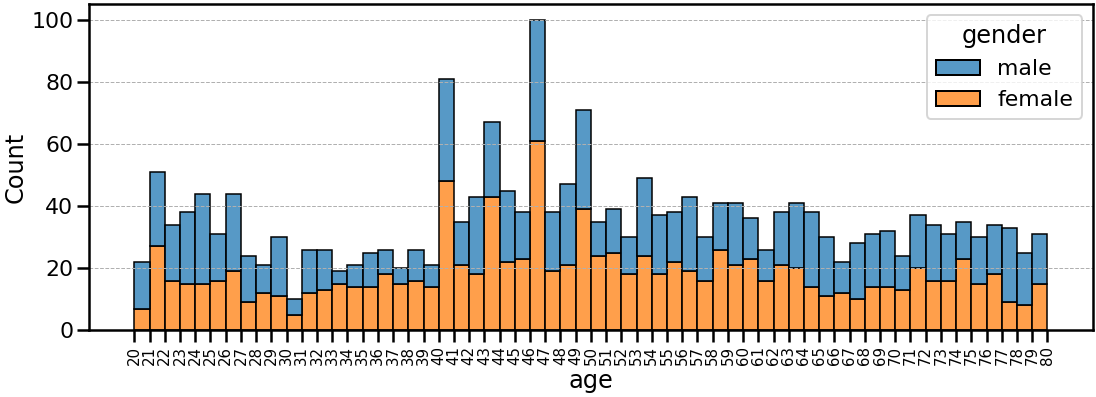

Figure S1: Age distribution of the sample dataset by gender

*Sigmoid fitting analysis,* fitting a sigmoid curve to the first derivative of the average β values, between 20 to 80 years, with 1year intervals for each of our non- stationary CpGs sites.

The function defined for optimization:

Variables (Scaling parameters):

1) S = responsible for scaling the output range from $[0,1]$ to $[0,S]$
2) B = adds bias to the output and changes its range from $[0,S]$ to $[B,S+B]$
3) K = responsible for scaling the input, which remains in $(-\infty,\infty)$
4) $\lambda$ = the point in the middle of the Sigmoid, i.e. the point where Sigmoid should originally output the value 0.5

The Model:

$$F\left( x \right)=\frac{S}{\left( 1+e^{-K*\left( x-\lambda\right)} \right)+B}$$

We used the least squares method to fit the model and find the optimal parameters ($S,B,K,\lambda$) for each one of the 67,882 processed and smoothed curves.

Probes that failed to converge in 5,000 iterations are discarded.

Table S2: 19 parameters derived from the piecewise regression and sigmoid models used for the establishment of the decision tree model utilized for distinguishing valid ESP probes from invalid probes

| Feature | Importance | Feature Explanation |
| --- | --- | --- |
| piecewise_-log(mse) | 0.174587 | The –log10 of the mean squared error in piecewise fit to signal |
| sigmoid_-log(mse) | 0.132899 | The –log10 of the mean squared error in sigmoid curve fit to signal derivative |
| peicewise_r2 | 0.113734 | The R^2 score of piecewise fit to signal |
| sigmoid_r2 | 0.10615 | The R^2 score of sigmoid curve fit to signal derivative |
| transition time | 0.061759 | The number of years between the two slopes in sigmoid model |
| sigmoid_pw3 _stop | 0.04587 | The year at which the second slope starts |
| sigmoid_fit_S | 0.030902 | The S parameter of the sigmoid model |
| peicewise_slope_B | 0.03042 | The B parameter in the sigmoid model |
| sigmoid_fit_$\lambda$ | 0.026238 | The middle point of the sigmoid curve |
| l_slope | 0.020467 | The linear regression slope |
| sigmoid_pw3 _start | 0.019983 | The End of the First slope in the sigmoid function |
| sigmoid_fit_K | 0.019592 | The K parameter in the sigmoid function |
| peicewise_sp | 0.018034 | The elbow point at the piecewise function |
| peicewise_delta | 0.015651 | The difference between the two slope a and slope b in piecewise fit |
| l_slope_-log(mse) | 0.013921 | The –log10 of the mean squared error in linear regression fit to signal |
| sigmoid_fit_b | 0.012704 | The second slope in the piecewise fit |
| l_slope_r2 | 0.011659 | The R^2 score of the linear regression fit |
| peicewise_slope_a | 0.011383 | The first slope in the piecewise fit |
| chr | 0.008697 | The chromosome on which the probe is located |

**Distribution of the β values on the type III ESPs**

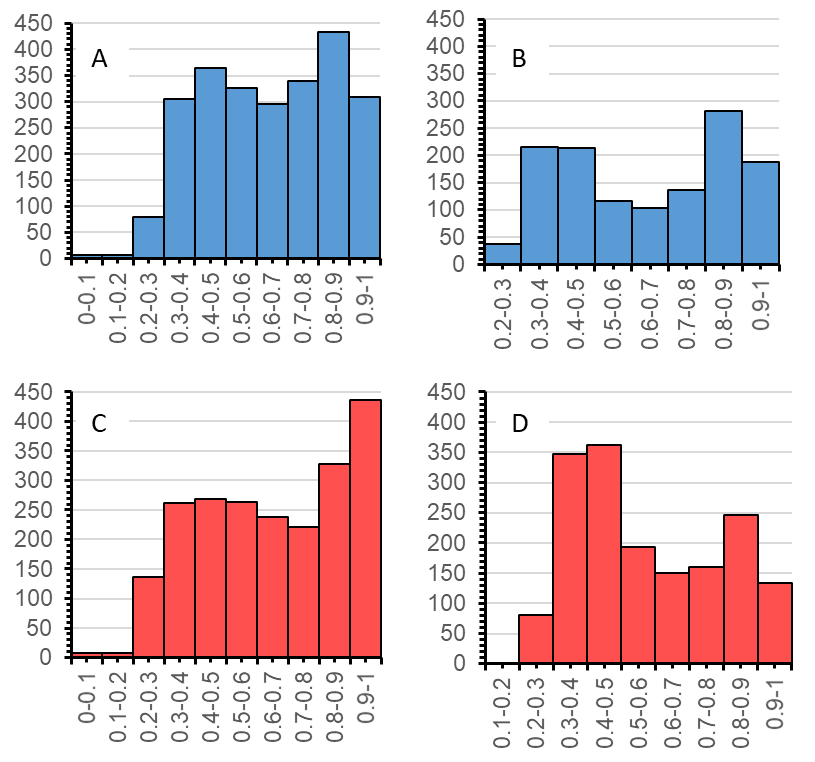

Figure S2: Distribution of β values in type III ESPs. A, B) distribution of β values in men, C, D) distribution of β values in women. A and C are for ESPs with a positive overall trend and B and D are for SP CpG with a negative overall trend.
